# Multichannel Sleep Spindle Detection using Sparse Low-Rank Optimization

**DOI:** 10.1101/104414

**Authors:** Ankit Parekh, Ivan W. Selesnick, Ricardo S. Osorio, Andrew W. Varga, David M. Rapoport, Indu Ayappa

**Author notes:** Corresponding author. Email address Source Code available at https://github.com/aparek/mcsleep.git.

## Abstract

**Background:** We propose a multichannel spindle detection method that detects global and local spindle activity across all channels of scalp EEG in a single run

**New Method:** Using a non-linear signal model, which assumes the multichannel EEG to be a sum of a transient component and an oscillatory component, we propose a multichannel transient separation algorithm. Consecutive overlapping blocks of the multichannel oscillatory component are assumed to be low-rank whereas the transient component is assumed to be piecewise constant with a zero baseline. The estimated multichannel oscillatory component is used in conjunction with a bandpass filter and the Teager operator for detecting sleep spindles

**Results and comparison with other methods:** Several examples are shown to illustrate the utility of the proposed method in detecting global and local spindle activity. The proposed method is applied to two publicly available databases and compared with 7 existing single-channel automated detectors. F_1_ scores for the proposed spindle detection method averaged 0.66 (0.02) and 0.62 (0.06) for the two databases, respectively. For an overnight 6 channel EEG signal, the proposed algorithm takes about 4 minutes to detect sleep spindles simultaneously across all channels with a single setting of corresponding algorithmic parameters

**Conclusions:** The proposed method aims to mimic and utilize, for better spindle detection, a particular human expert behavior where the decision to mark a spindle event may be subconsciously influenced by the presence of a spindle in EEG channels other than the central channel visible on a digital screen

## 1. Introduction

Sleep spindles are short rhythmic oscillations visible on an electroencephalograph (EEG) during non-rapid eye movement (NREM) sleep. The center frequency of sleep spindles is between 11 and 16 Hz [58]. The duration of sleep spindles is defined to be at least 0.5 seconds, with some studies imposing an upper limit on their duration to 3 seconds [67]. Sleep spindles reflect a heritable set of traits which is implicated in both sleep regulation and normal cognitive functioning [38]. Recent studies have linked spindle density (number of spindles per minute), duration and amplitude of spindles, and peak frequency of spindles to memory consolidation during sleep [31, 15], cognition in schizophrenia patients [38, 66], brain dysfunction in obstructive sleep apnea [14] and biomarkers for Alzheimer’s disease [69]. As a result, understanding the characteristics of sleep spindles is a key in studying their relation to several neuropsychiatric diseases.

Traditionally, sleep spindles are annotated in clinics using visual heuristics: number of peaks or bumps of the EEG signal are counted within a specified time window. This process is not only subjective and time-consuming, but also prone to errors. Moreover, visual inspection underscores the fine details of putated spindles [51]. In order to reduce the subjectivity of visual detection, it is not uncommon for studies to utilize more than one expert for detecting spindles. However, in several cases this leads to a high variability in inter-scorer agreement. The Cohen’s *κ* coefficient for inter-rater agreement in manual scoring usually ranges between 0.46 and 0.89 [60, 42]. As such, the presence of reliable automated spindle detectors may not only reduce the scoring variability [71] but may also aid in complex longitudinal studies that involve studying global or local sleep spindle dynamics [51, 21, 43].

Broadly categorized, there exist two-types of automated sleep spindle detectors for single channel EEG: filtering based and non-linear signal decomposition based. Filtering based approaches vary from basic methods, which utilize a bandpass filter with constant or adaptive thresholds, to advanced methods that use time-frequency information along with bandpass filtering. Most of the filtering based methods involve pre-processing of the desired channel of the EEG (usually a central channel) for artifact removal [35]. One of the first automated detectors to be proposed used a bandpass filter in conjunction with an amplitude threshold [56]. This idea is still the basis of a majority of the bandpass filtering-based automated detectors [68, 22, 39, 30, 16, 32]. Advanced methods utilizing time-frequency information either use a wavelet transform [36, 2, 4, 29, 63, 3] or a short-time Fourier transform (STFT) [20, 46, 24] with adaptive thresholding to detect spindles. Several machine-learning based spindle detectors and sleep staging algorithms have also been proposed for single channel EEG [1, 34].

Non-linear signal decomposition based methods [49, 50, 36, 26] attempt to separate the non-rhythmic transients or artifacts from sinusoidal spindle-like oscillations in the single channel sleep EEG. These methods make use of the differing morphological aspects [59] of the transients and spindles to overcome the drawbacks of filtering and Fast Fourier Transform (FFT) based techniques [52]. As an another example, Gilles et. al considered the removal of ballistocardiogram (BCG) artifacts from EEG using lowrank and sparse decomposition [33]. In addition to these morphological component analysis (MCA) based methods, independent component analysis (ICA) and principal component analysis (PCA) have also been used to detect spindles for single channel EEG [5]. However, note that ICA assumes linearity and stability of the mixing process along with statistical independence of input sources [27].

### 1.1. Motivation

Automated spindle detectors that consider only a single channel are blind to the presence of spindles in other recorded channels. Such a spindle detection mechanism may not be in concordance with the way spindles are annotated visually. The American Academy of Sleep Medicine (AASM) manual for scoring of sleep and associated events recommends using F4, C4 and O2 channels (or alternatively Fz, Cz and C4) of the recorded EEG with F3, C3 and O1 as backup channels [58]. As such, while annotating sleep events rarely does an expert view a single channel of the EEG in isolation to the other channels. This is certainly the case for studies either looking to characterize individual global sleep spindle density [9] or tracking the propagation of spindles overnight [51, 65]. As a result, it may be possible that the presence of spindles in channels other than the channel of interest subliminally influences the experts’ decision of marking an event as a spindle.

As an example, consider the 3-channel EEG shown in Fig. 1. The experts visually annotated the presence of a spindle at approx. 26 seconds. While it is suggested that only the central channel was used for annotating spindles [22], it can be seen that the spindle at approx. 26 seconds is also present in the frontal and the occipital channels, though with different amplitudes. As such it is highly likely that the decision by a human to mark the presence of a spindle at approx. 26 seconds in the central channel is reinforced by its presence in other channels if they are viewed together on a digital screen. Similar behavior can be seen in the case of the EEG excerpt in Fig. 2 where the experts annotated a spindle at approx. 29.5 seconds. While the degree to which the decision of marking a spindle was influenced by its presence in other channels (if it occurred simultaneously in more than one channel) is an open question out of the scope of this paper, utilizing it can certainly aid in a better design of the automated spindle detectors.

**Figure 1:**
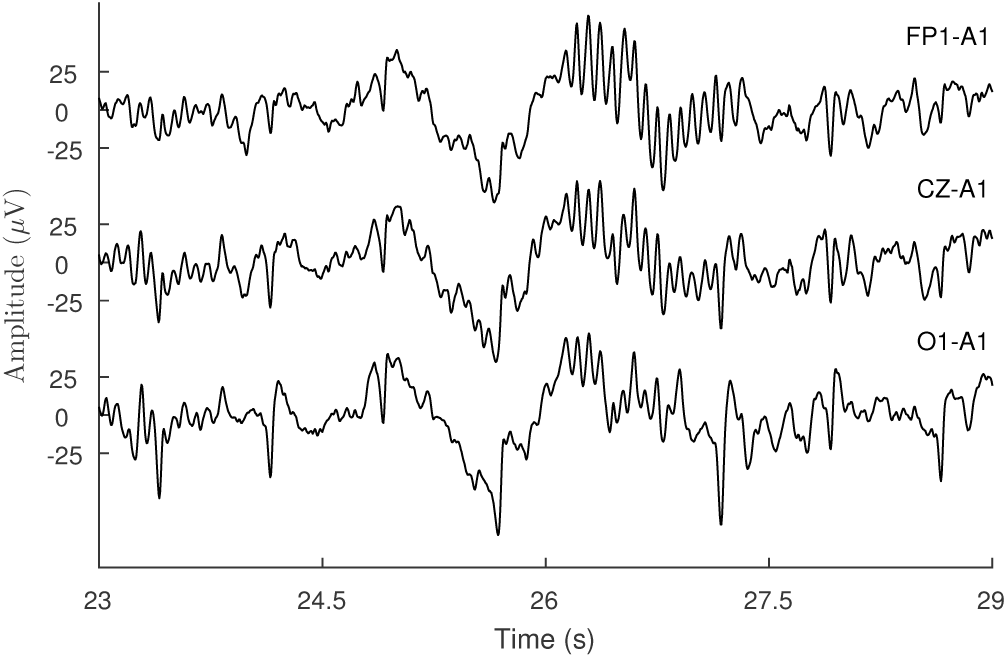
A multichannel EEG with one frontal, one central and one occipital channel. Expert annotated sleep spindle at 26 seconds has different amplitude in different channels.

**Figure 2:**
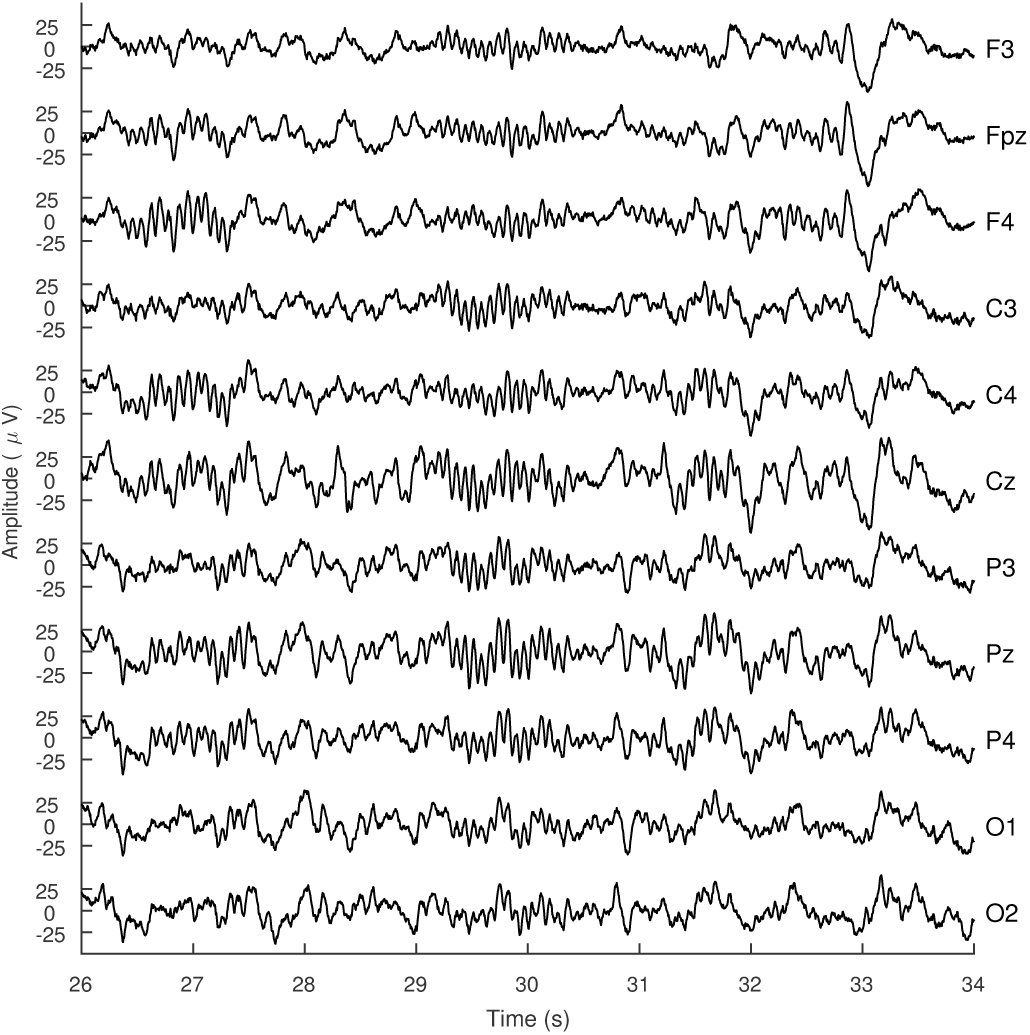
An example of a multichannel scalp EEG from the MASS database [45] (SS2, PSG1, epoch 701). The spindle at approx. 27 seconds appears predominantly in the right channels. The visual annotation of spindle at approx. 29 seconds is reinforced by the presence of spindle in other channels.

Another motivation for considering multichannel EEG for studying spindle activity comes from the fact that while single channel detectors may be used to study global spindle activity, their usage comes at a price. Since the amplitude of spindles vary in each channel (see for example Fig. 1), amplitude-based thresholds used by automated detectors need to be tuned separately for each channel, adding to the overall computational complexity. Additionally, the CPU time is multiplied by the number of channels recorded. While this additional computing time may not be significant for the case of basic filtering-with-thresholding methods, it is certainly significant for advanced methods that utilize either time-frequency information or non-linear signal decomposition.

Classifying spindles as either global or local [15, 12] is difficult using single channel based methods: spindles that appear on the right channels (F4, C4, and O2 channels in Fig. 2) may be entirely missed by detectors using the left channels (F3, C3 and O1) or vice-versa, which is the case with most detectors [67]. In fact, most detectors utilize at the most two channels [44, 52] possibly from the same hemisphere (left or right). Thus, for a single channel method to effectively detect global spindle activity it has to be run on channels belonging to the left, right hemispheres and also the midline channels. As an example, consider the spindle at approx. 26.5 seconds in the EEG shown in Fig. 2. This spindle appears prominent in the right channel, while appearing in other channels (such as Pz and Cz) with significantly lower amplitude. As a result, without careful parameter tuning an automated detector may not be able to properly detect the spindles. Furthermore, combining the detections to a single binary decision vector is also challenging.

### 1.2. Contribution

The contributions of this paper are two-fold. First, we propose a multichannel transient separation method which decomposes the input multichannel EEG as the sum of a transient and an oscillatory component. Second, we utilize the estimated oscillatory component to detect spindles by using an envelope of the bandpass filtered oscillatory component. Combined, the two contributions of this paper aim to detect spindles across all channels of scalp EEG in a single run with a single parameter-tuning (i.e., the parameter-tuning is independent of the number of recorded EEG channels).

The transient component, in the proposed non-linear signal model, is modeled as a sparse piecewise constant signal, whereas the oscillatory component is assumed to exhibit block-similarity, i.e., the blocks of the multichannel oscillatory component are low-rank arrays. We estimate the two components of the proposed non-linear signal model by formulating an optimization problem posed as the minimization of a convex objective function. We derive a fast matrix-inverse-free algorithm to obtain the solution of the proposed optimization problem. Using an envelope of the bandpass filtered oscillatory component we detect sleep spindles globally i.e., across all scalp channels of the recorded multichannel EEG.

The bandpass filter used for the detection of spindles is excited by the presence of transients such as BCG, electrocardiogram (ECG) and other non-rhythmic waveforms. By separating the transients from the input multichannel EEG, the proposed method avoids the unnecessary excitation of the bandpass filter. Thus, the spindle activity appears more prominently than if the bandpass filter was directly applied to the input EEG. Moreover, due to the effective attenuation of non-oscillatory transients, a low-order bandpass filter suffices for sleep spindle detection [50, 49].

### 1.3. Relation to prior work

Several studies have pointed to the benefit of separating the transients and oscillations sleep spindle detection [50, 49, 65], though few have advocated the use of multichannel EEG [6]. Non-linear signal models for the input EEG have been used to detect sleep spindles directly for single channel [50, 49] and indirectly for multichannel EEG [6]. An ICA based approach was studied for the detection of spindles from multichannel EEG in [54]. For sleep-staging and classification a machine learning approach utilizing multichannel EEG was proposed in [57]. A matching pursuit (MP) based decomposition method was proposed for multichannel EEG to relax the assumptions of ICA [27]. The MP based method attempts to represent the multichannel EEG as a linear combination of atoms or coefficients with respect to a chosen basis. Estimating the atoms, by solving an inverse problem, can enable detection of sleep spindles with user chosen parameters [27]. Similar to the ICA approach, a multichannel Matching Pursuit based method was also proposed for the decomposition of the multichannel EEG signal [61]. An MP-based algorithm using singular value decomposition (SVD) was shown to be able to efficiently learn the different oscillatory waveforms in multichannel EEG [11].

### 1.4. Organization

The rest of the paper is organized as follows. In Section 2 we detail the notation used throughout the paper and introduce the block low-rank operator. In Section 3 we propose a non-linear signal model for the EEG and show how to estimate its components. We formulate and derive the transient separation algorithm and show how to use the estimated components for detecting sleep spindles globally. In Section 4 we illustrate the proposed transient separation and spindle detection methods and show how to set the parameters for the same. In Section 5 we illustrate the proposed method on publicly available annotated single channel spindle databases and finally conclude in Section 6.

## 2. Preliminaries

### 2.1. Notation

We denote vectors and matrices by lower and upper case letters respectively. An n-point signal *y* is represented by the vector

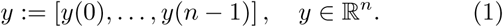

We represent the multichannel signal *X* ∈ ℝ^*k×n*^, with *k* channels as

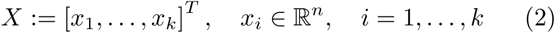

where [*·*]^*T*^ represents the transpose. The *ℓ*_1_ and *ℓ*_2_ norm of the vector *y* are defined as

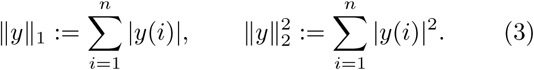

The nuclear norm of the matrix *X ∈* ℝ^*m×n*^ is defined as

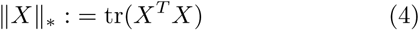

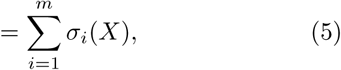

where tr(*·*) represents the trace and *σ*_*i*_(*X*) is the *i*^th^ singular value of *X*.

We define the matrix *D* ∈ ℝ^(*n−*^1^)*×n*^ as

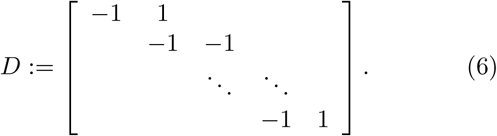

Using the matrix *D*, the first-order difference of a discrete signal *y ∈* ℝ^*n*^ is given by *Dy*. The soft-threshold function [25] for *λ >* 0 is defined as

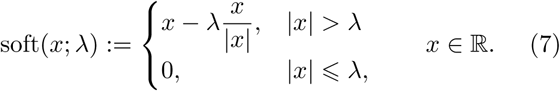

Note that the soft-threshold function in (7) is applied element wise to a vector with threshold *λ >* 0.

The Teager-Kaiser energy operator for a discrete-time signal *y* denoted by *T* (*·*) is defined as

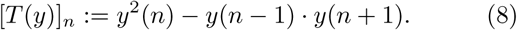

Note that, unless stated otherwise, applying the Teager operator to a multichannel signal *X* implies that the Teager operator is applied to the channel mean of *X*. The Teager operator has been commonly used to obtain an envelop of the bandpass filtered signal for spindle detection [46, 3, 29, 20, 67].

### 2.2. Block Low-Rank Operator

In order to extract the rhythmic oscillations (i.e., sleep spindles) from the multichannel EEG, we propose the following sparse optimization framework. Consider a segment of a sample multichannel EEG, as shown in Fig. 3(a) with three blocks highlighted (each of 1 second in length). Expert annotated spindle at approx. 26 seconds overlaps the second multichannel block i.e., Block 2. Figure 3(b) shows the singular values of each of the three blocks. It can be seen in Fig. 3(b) that the Block 2 has larger (in magnitude) singular values. As such, if one were to approximate each of the three blocks with their low-rank approximations^1^, the low-rank approximation for Block 2 would still contain the spindle-like oscillations. Figure 3(c) shows the multichannel EEG from Fig. 3(a) with the blocks replaced by their corresponding rank-one approximation. Furthermore, in the absence of non-spindle like transients the sum of the singular values of all blocks in the multichannel EEG will be approximately equal to the sum of singular values of the spindle-containing blocks. As a result, in order to extract the spindles, it suffices to obtain a low-rank representation for multichannel EEG, once non-rhythmic transients are separated.

**Figure 3:**
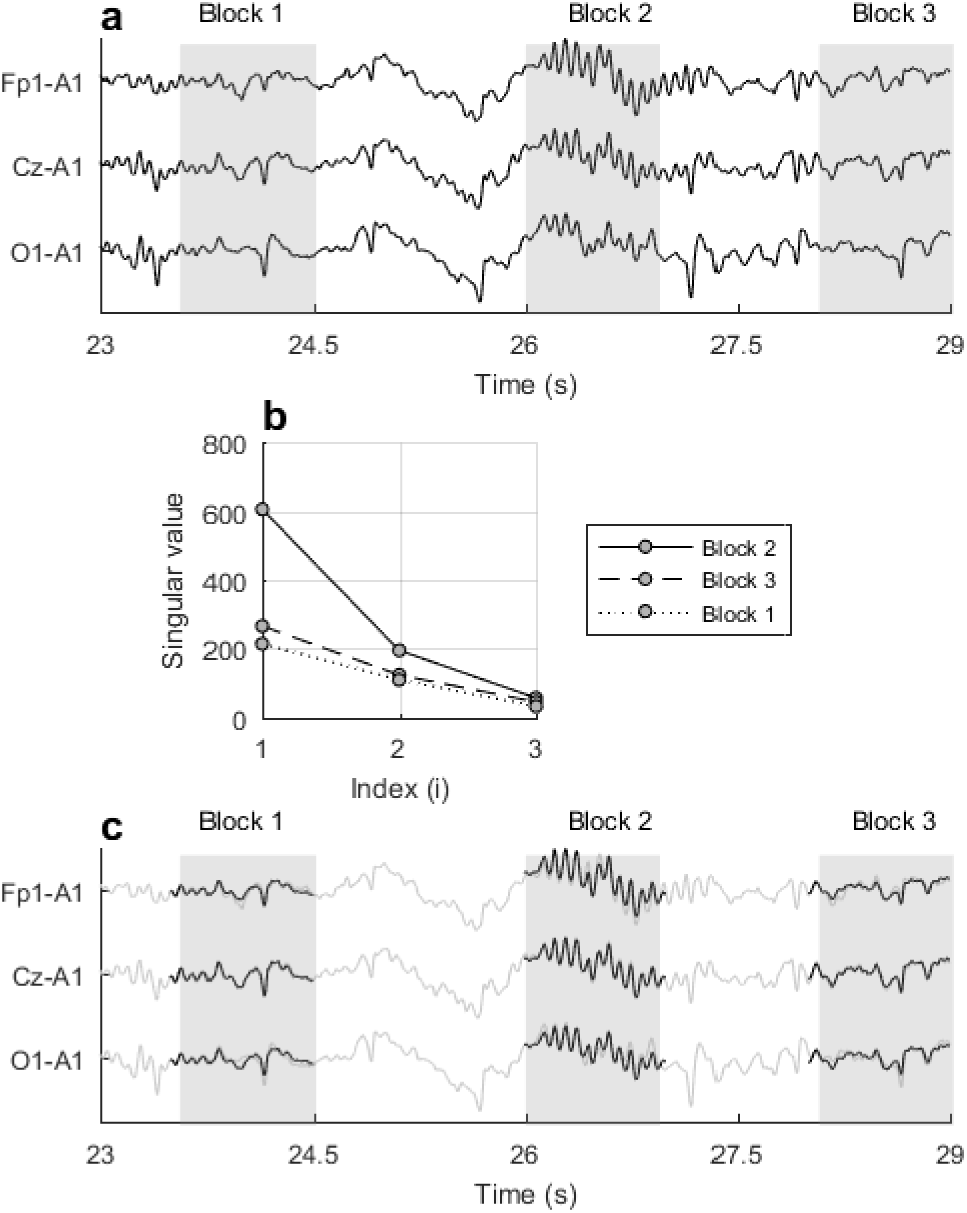
(a) Sample multichannel EEG with multichannel blocks that contain transients and spindles are highlighted (length of each block is 1 second). (b) Singular values of each of the highlighted blocks in (a) are shown. (c) Sample multichannel EEG with the highlighted blocks replaced by their rank-one approximation. Shown in the background is the multichannel EEG from (a).

**Figure 4:**
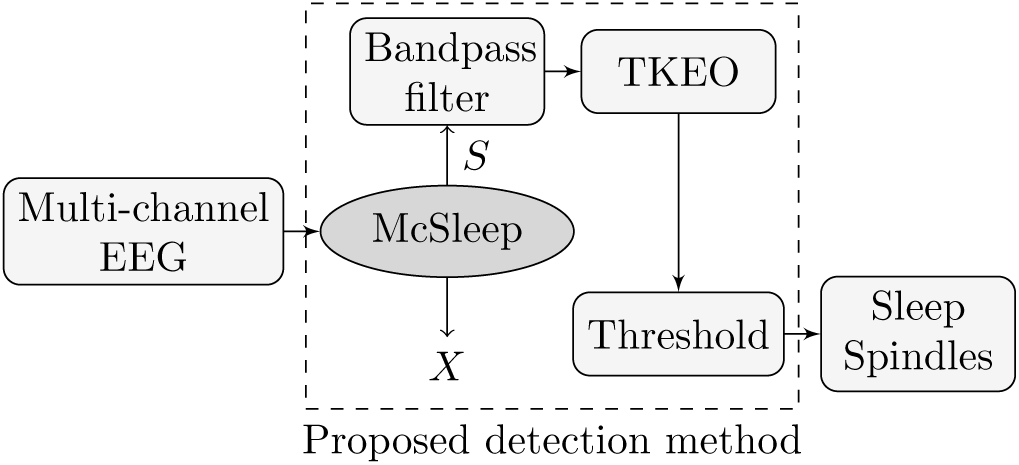
Proposed detection method for multichannel EEG using McSleep as a multichannel transient separation algorithm.

We define the operator Φ: ℝ^*k×n*^ → ℝ^*k×l×m*^, which extracts *m* blocks, each of an even length *l*, from the *k−*channel input signal as

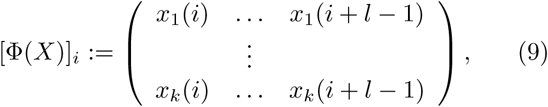

for *i* = 1, …, *m*. The operator Φ can be defined with a certain overlap between consecutive blocks. Further, Let the adjoint operator be denoted by Φ^*T*^: ℝ^*k×l×m*^ *→* ℝ^*k×n*^. The adjoint operator forms the *k*−channel signal by aggregating the *m* blocks, where by aggregating we mean that the blocks are added in an overlap-add way. Note that in the case of distinct blocks, i.e., no overlap between the blocks, the operator Φ is orthogonal (Φ^*T*^ Φ = *I*).

In this paper, we use the operator Φ with 50% overlap between blocks of 1 second in length, implemented to obtain perfect reconstruction. As an example, for an EEG signal sampled at 256 Hz, the block length is fixed at 256 samples. In order to perfectly reconstruct the input signal *X* ϵ ℝ^*k×n*^ from Φ(*X*), we use a diagonal weight matrix *W* ϵ ℝ^*n×n*^. Since the blocks are aggregated in an overlap-add way, the samples of signal *X* that are contained in the overlap occur twice in the signal formed using the adjoint operator Φ^*T*^. As a result, appropriately weighting the samples can lead to perfect reconstruction^2^.

An example will help clarify the proposed block lowrank operator. Consider the single channel signal *X* = [*x*_1_*, x*_2_*, x*_3_]. Using a block length of 2 samples with 50% overlap leads to

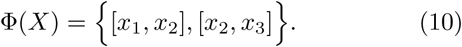

Reconstructing the signal from the individual blocks by overlapping and adding we get

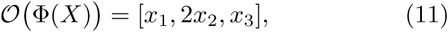

where *O*(*·*) defines the overlap-add operator. Note that by ‘overlap-add’ we imply that the individual blocks of size *k × l* are overlapped and added to construct a multichannel signal of size *k × n*. In order to achieve perfect reconstruction, i.e., Φ^*T*^ (Φ(*X*)) = *X*, we use the weight matrix *W* ∈ ℝ^3*×*3^, given by

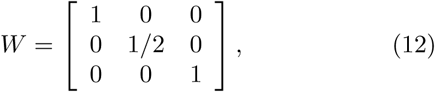

and define

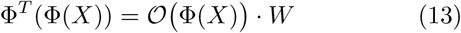

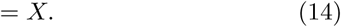

Note that the weight matrix *W* associated with the operator Φ with 50% overlap can be pre-computed based on the input signal length and the user chosen block length. In particular, for an input signal *X* ∈ ℝ^*k×n*^, *n >* 3, the diagonal weight matrix associated with the operator Φ with 50% overlap and an even block length of *l* is given by

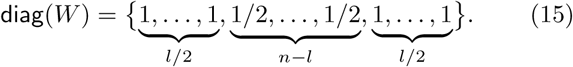

A suitable optimization problem for estimating the oscillatory component with a block low-rank structure is given by

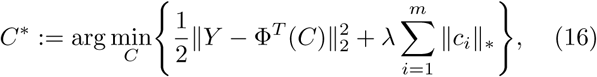

where *C* = [*c*_1_ …, *c*_*m*_], *c*_*i*_ ∈ ℝ^*k×l*^, *C\** ∈ ℝ^*k×l×m*^, and *λ >* 0 is the regularization parameter. Note that the optimization problem in (16) estimates the blocks *c*_*i*_ from which the multichannel signal *S* can be calculated using Φ^*T*^ (i.e., *S* = Φ^*T*^ (*C*^*\**^)). The optimization problem in (16) is a sum of convex functions (nuclear norm) and a strictly convex function (*ℓ*_2_ norm squared) and hence is a convex optimization problem. As a result, well developed principles of convex optimization can be leveraged to obtain a global minimum [48].

The solution to the optimization problem in (16) can be obtained using the iterative shrinkage/thresholding algorithm (ISTA) [7] and its variants. The ISTA algorithm, for the optimization problem in (16), entails soft-thresholding the singular values of each block of the multichannel signal *Y*. For an overnight multichannel EEG signal, roughly 30000 blocks of length 1 second are obtained using the operator Φ and as such the ISTA algorithm involves computing 30000 singular value decompositions (SVD). However, since the number of channels is much less than the length of the block, we need compute only the first *k* (number of channels) singular values and their corresponding left and right singular vectors. As an example, for the multichannel EEG signal shown in Fig. 3 or for the one in Fig. 2 it suffices to compute only the first 3 or 11 singular values respectively.

## 3. Sleep Spindle Detection for Multichannel EEG

### 3.1. Non-linear signal model

We propose the following non-linear signal model for the multichannel EEG denoted by *Y*:

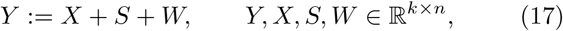

where *X* represents the transient component, *S* represents the oscillatory component and *W* represents additive white Gaussian noise (AWGN) (i.e., *W ∼ 𝒩*(0, *σ*)). We assume that the transient component *X* is sparse and piece-wise constant and the blocks of the oscillatory component *S* are low-rank as described in Sec. 2.2.

The signal model presented in this paper contains certain similarities to the one presented in [50], in particular, the transient component is modeled in a similar way. Moreover, in both models, sparsity of the block structure of the spindle component is exploited. Although, the properties of the spindle component presented in this paper are different, the overall theme of non-linear signal models presented in this paper, in [50] and in [49] is similar: the input EEG signal is modeled as a sum of transient and oscillatory components.

### 3.2. Estimating Transient and Oscillatory Components

In order to detect spindles, we first estimate the transient and the oscillatory components in the proposed signal model (17) from the recorded multichannel EEG. To this end, we utilize a sparse optimization framework and propose to solve the following objective function

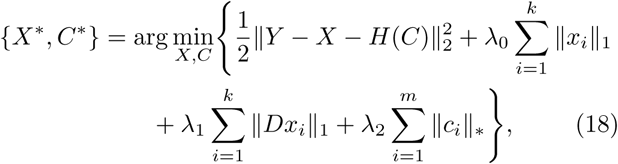

where *X* = [*x, …, x*], *C* = [*c, …, c*], *c* ∈ ℝ^*k×l*^ and *λ*_*i*_ *>* 0 are the regularization parameters. We let *H* = Φ^*T*^, where Φ is the block low-rank operator (see Sec. 2.2). Recall that *D* is the first-order difference matrix, as defined in (6), and *C* is the coefficient array obtained using the operator Φ as defined in (9).

The proposed objective function seeks the optimal solution *X\** which is sparse and piecewise constant. The ℓ_1_ norm on *X* penalizes non-sparse solutions and the *ℓ*_1_ norm on *Dx*_*i*_, for *i* = 1*, …, k*, penalizes non piecewise constant solutions. These two penalties combined are generally termed as the ‘fused-lasso’ penalty [62, 47] and have been shown to model the transient component [50] with relative accuracy.

The nuclear norm on each of the coefficients *c*_*i*_ in (18) penalizes solutions *C*^*\**^ that do not exhibit the block lowrank property as described in Sec. 2.2. Using the solution *C*^*\**^ from the optimization problem in (18), we estimate the oscillatory component using the operator *H* = Φ^*T*^, i.e.,

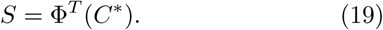

The estimate for the oscillatory component *S* can then used to detect sleep spindles.

### 3.3. Transient Separation Algorithm

We develop a fast iterative algorithm to obtain the optimal solution for *X*^*\**^ and *C*^*\**^ using the proposed objective function (18). Note that the objective function proposed is convex and hence well developed theory of convex optimization algorithms can be leveraged to obtain the optimal solution. We apply Douglas-Rachford splitting [18] to solve (18), which results in an instance of the alternating direction method of multipliers (ADMM) method. The convergence of the iterative ADMM algorithm is guaranteed for the proposed objective function (18) under suitable assumptions [28, 10].

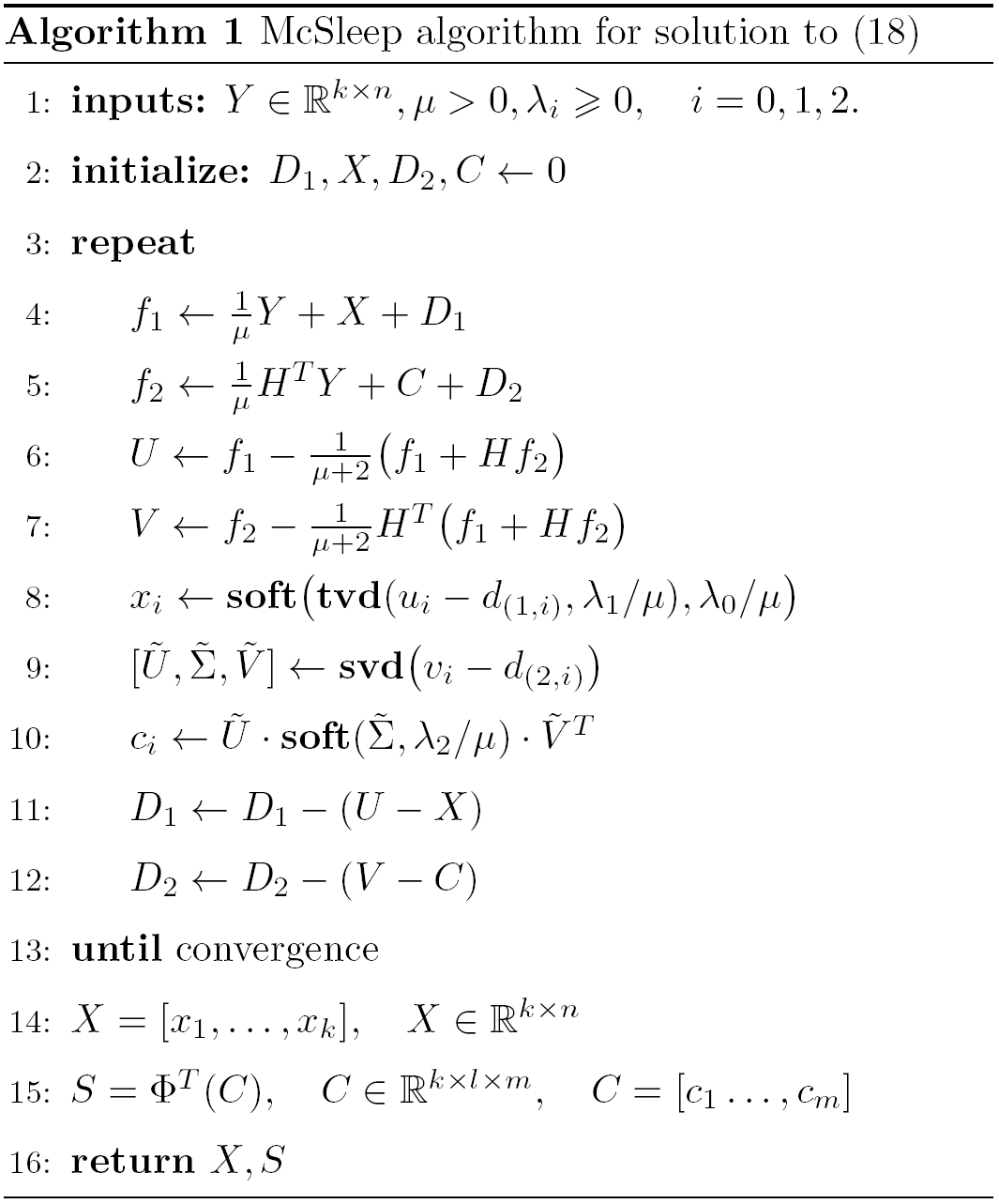

We write the objective function (18) using variable-splitting as

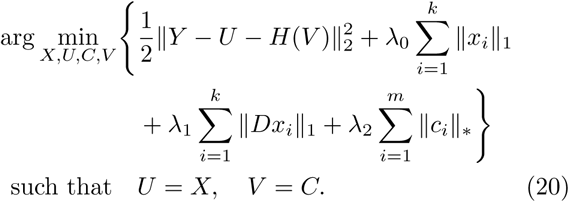

Using the scaled augmented Lagrangian, minimizing (20) results in solving the following three sub-problems:

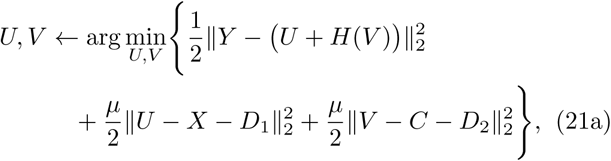

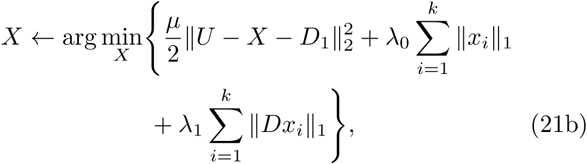

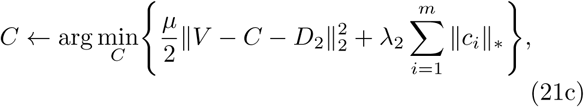

where *µ >* 0 is the Lagrangian step-size parameter.

The first term in sub-problem (21b) can be written as the energy over each channel of *U, X* and *D*_1_, i.e.,

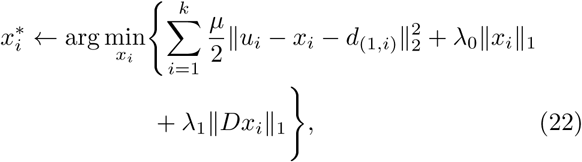

with 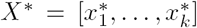. The terms *u*_*i*_*, x*_*i*_ and *d*_(1*,i*)_, for *i* = 1*, …, k* represent the *k−*channels of *U, X* and *D* respectively. The solution to (22), for each 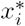, is readily obtained by applying the fused-lasso method [62] to each channel of the underlying signal, i.e.,

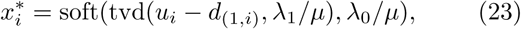

where *u*_*i*_ and *d*_(1*,i*)_ are the *i*^th^ channel of *U* and *D*_1_ respectively. Note that tvd(·) represents the solution of total variation denoising method [55] obtained using a fast solver [19] and soft(·) represents the soft-thresholding function (7).

We write the sub-problem (21c) as

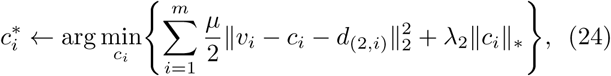

where *v*_*i*_*, c*_*i*_ and *d*_(2*,i*)_ are the *i*^th^ channel of *V, C* and *D*_2_ respectively. The solution to (24) is obtained using the singular value thresholding (SVT) algorithm [13], i.e.,

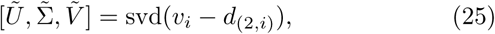

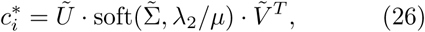

where svd(*·*) represents the singular value decomposition. The SVT algorithm computes the singular values of the input matrix and thresholds them using the soft-threshold function [13].

The objective function in the sub-problem (21a) can be solved exactly using a suitable substitution via the least squares method. Note that the objective function in the sub-problem (21a) is similar to objective function (20a) in [50], and hence a similar derivation can be used for (21a) in this paper. We detail the derivation in Appendix 9.1. The iterative algorithm for (18) is listed in Algorithm 1 and the MATLAB code is made available online^3^.

Figure 5 shows the estimated transient and oscillatory components for a three channel EEG (FP1-A1, CZ-A1, O1-A1) from the DREAMS Database^4^. It can be seen in Fig. 5 that the spindles in the three EEG channels are captured by their respective oscillatory components, whereas the non-oscillatory waveforms are captured by the transient components. Also shown in Fig. 6 is the separation of transients and oscillations for a sample 6-channel EEG from the Montreal Archive of Sleep Studies (MASS) database [45]. The regularization parameters for the preceding examples were found empirically.

**Figure 5:**
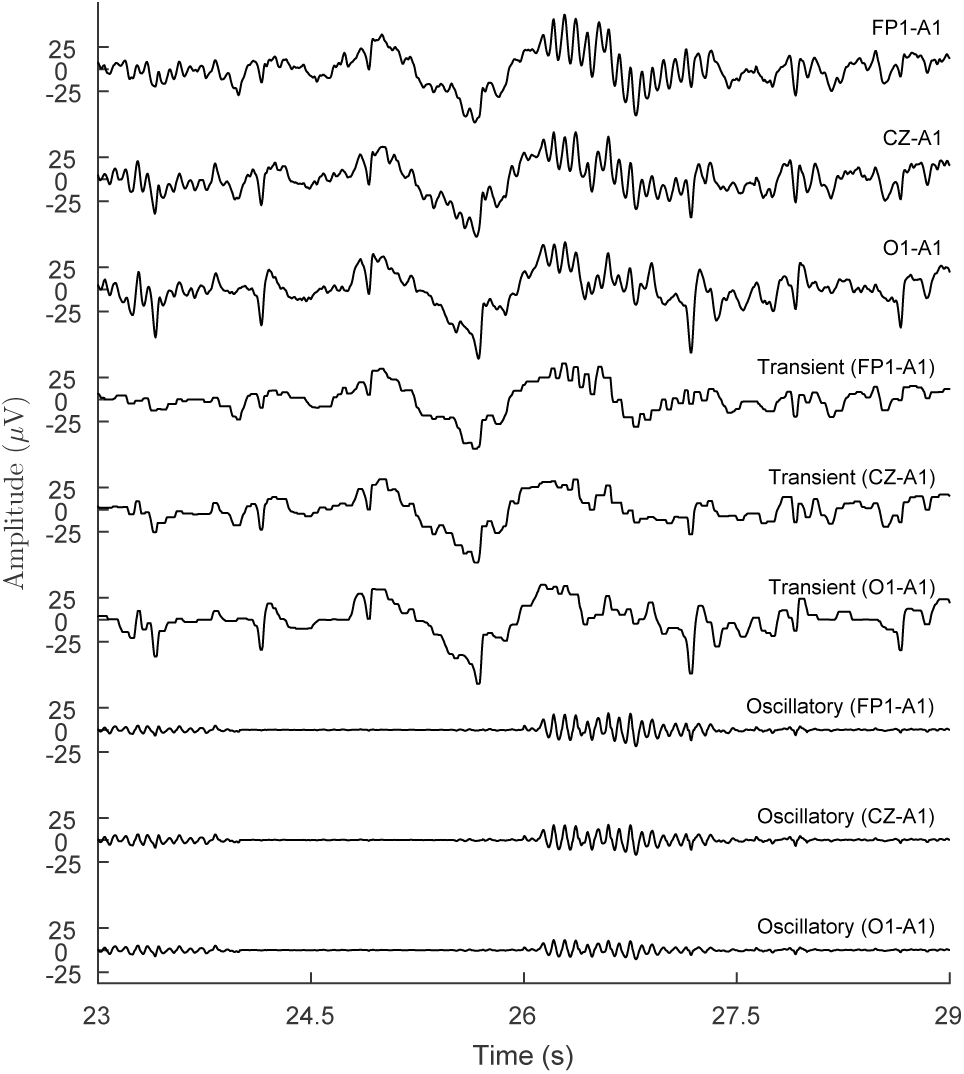
Separation of transients and oscillations using the proposed objective function in (18) for an example EEG segment from DREAMS Database [23]. The transient component is modeled as the sum of a low-frequency signal and a sparse piecewise constant signal.

**Figure 6:**
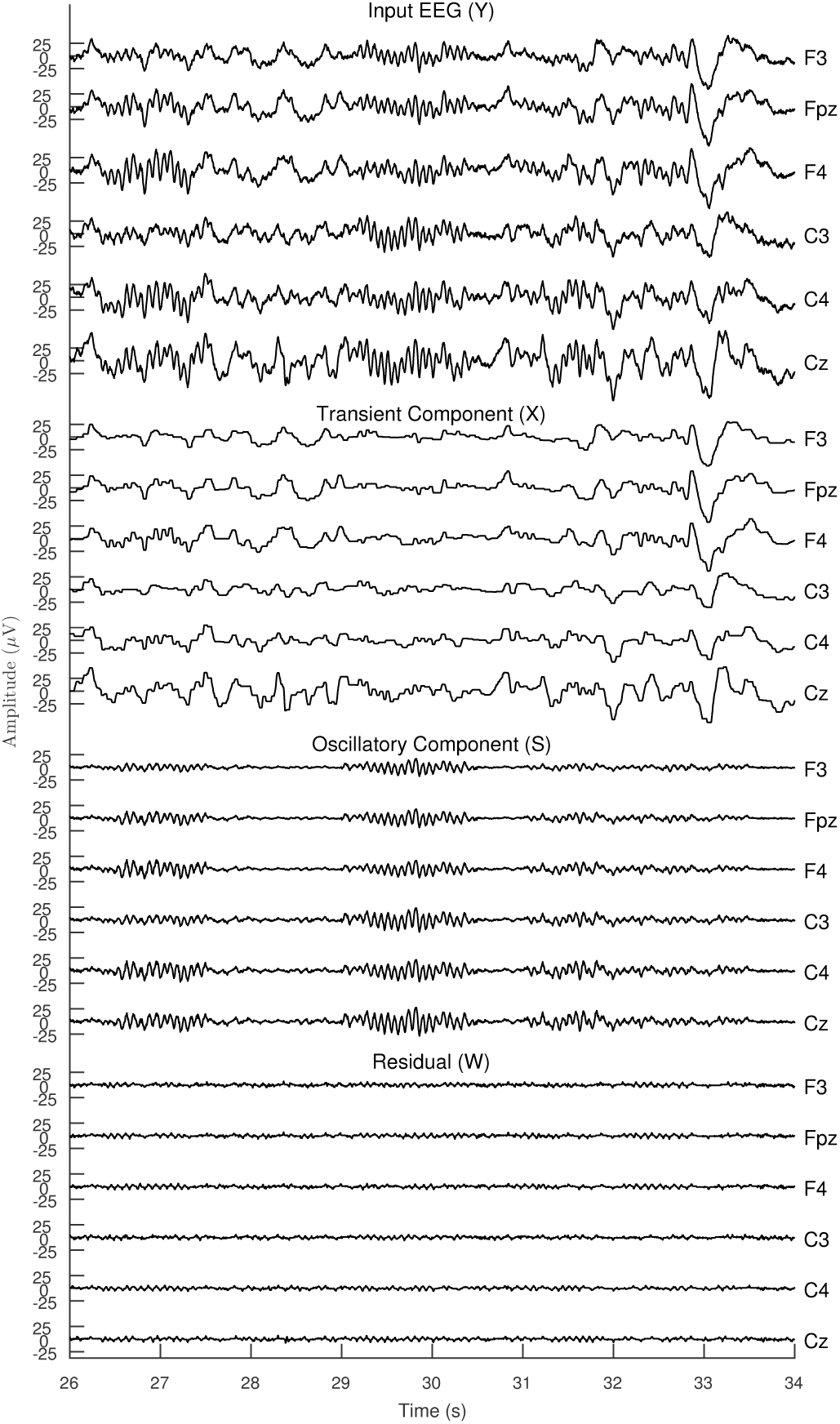
Decomposition of a 6-channel EEG excerpt (*Y*) into its transient (*X*) and oscillatory (*S*) components. Also shown is the the residual *W*, such that (*Y* = *X* + *S* + *W*). The EEG excerpt is same as the one in Fig. 2. N.B. For clarity only first 6 channels are shown.

### 3.4. Detection of Spindles Post Separation of Transients

We use the estimated multi-channel oscillatory component to detect the sleep spindles. In order to suppress non-spindle like waveforms captured by the oscillatory component, we use a 4^th^ order Butterworth bandpass filter with a user-specified passband. Specifically the bandpass filter is applied to each channel of the estimated oscillatory component. We denote the bandpass filtered oscillatory components as BPF(*S*), where *S* is the oscillatory component.

The usage of the proposed transient-separation algorithm, described in the preceding subsection, allows for the oscillatory activity in the EEG to appear prominently. As a result, post separation of the transients the detection of sleep spindles becomes relatively simpler. We use the Teager Operator, as defined in Sec. 2, to construct an envelope of the oscillatory activity and consequently detect spindles. The Teager operator denoted by *T* (·), is applied to the channel mean of the multichannel bandpass filtered oscillatory component (BPF(*S*)) to detect the global spindle activity. Using a constant threshold, we define a binary signal bspindle(t) as

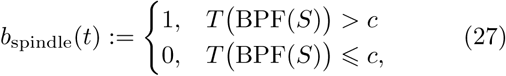

where 1 denotes a spindle present and 0 otherwise. Figure 4 summarizes the proposed multichannel sleep (McSleep) spindle detection method using the derived transient separation algorithm for sleep EEG.

## 4. Examples

We illustrate the proposed multichannel sleep spindle detection (McSleep) and compare it to other state-of-the-art automated single-channel based spindle detectors. To the best of the authors knowledge, an automated method for detecting spindles globally in a single run has not been studied before. Recall that the proposed method consists of two parts: separation of transients and oscillations from multichannel EEG and using bandpass filter with Teager operator to detect spindles. Hence, as discussed in Sec. 1.1, the proposed method can be applied in the following two scenarios. First, when studying global spindle activity the proposed method can be applied as described in the preceding subsection, i.e., by applying the Teager operator to the channel mean of the bandpass filtered oscillatory component. Second, if the user is interested in spindles only in a single channel then the Teager operator, postestimation of the oscillatory component, can be applied specifically to that channel. While the latter is a limitation of the proposed method in that only a single channel is considered post estimation of transients, the presence of multiple EEG channels can certainly aid in better separation of transients and oscillations for the input EEG. Before we illustrate an example for each of the two scenarios, we provide an overview of the parameters that need to be set for the proposed McSleep spindle detection method.

### 4.1. Parameters

The proposed spindle detection method requires the user to set several parameters which are either algorithm-specific or task-specific. Algorithm specific parameters are the regularization parameters *λ*_*i*_ ≥ 0, for *i* = 0, 1, 2., in (18) and the step-size *µ* for the scaled augmented Lagrangian. The regularization parameters influence the sparsity of their respective components. For example, a high value for *λ*_0_, relative to *λ*_1_ and *λ*_2_, enforces the transient component *X* to be sparse (i.e., with a baseline of zero). Similarly, a high value for *λ*_2_, relative to *λ*_0_ and *λ*_1_ results in the estimated oscillatory component to be of reduced rank. For a sufficiently high *λ*_2_, a rank-one approximation may be obtained.

The values for *λ*_0_, *λ*_1_ and *λ*_2_ were found empirically for the examples that follow. We find that the same *λ*_0_, *λ*_1_ and *λ*_2_ work well for different EEG signals having the same sampling frequency. Thus for a database that contains EEG signals sampled at the same frequency we can preset the values for *λ*_0_, *λ*_1_ and *λ*_2_. In case an EEG contains relatively more transients, the values of *λ*_0_ and *λ*_1_ may be increased proportionally. The step-size parameter *µ* on the other hand controls the rate of convergence for the proposed algorithm. Note that *µ* influences the speed at which the algorithm converges and not the solution to which it converges. Figure 7 shows the value of the proposed objective function (18) at each iteration for several values of *µ*. For the examples that follow and for the experiments in Sec. 5, we fix *µ* = 0.5. Note that setting the value of *µ* arbitrarily low or arbitrarily high may affect the convergence of the proposed transient separation algorithm.

**Figure 7:**
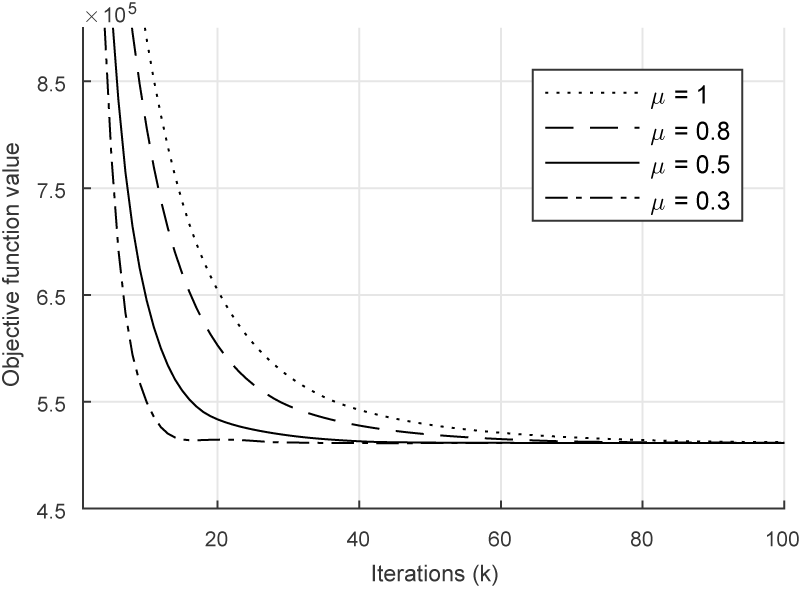
The value of objective function in (18), for each iteration *k*, is shown for several values of the step-size parameter *µ*.

The task-specific parameters are the block length for the block low-rank operator *H*, the overlap between consecutive blocks and the threshold *c* for the binary vector *b*_spindle_(*t*). The average duration of a sleep spindle is between 1 and 1.5 seconds [67, 4, 51]. As a result, we set the block length to be fixed at 1 second with a 50% overlap between consecutive blocks. The passband for the Butterworth filter used in this paper is in fact an additional parameter to be set when using the proposed method. We select the passband to be 11 Hz to 16 Hz based on spindle frequency range reported in [22] and the AASM manual [58]. However, increasing number of studies are reporting spindle frequencies to be in a variety of ranges, such as 11–15 Hz [41], 10–17 Hz [67]. As such, there seems to be a lack of consensus among the sleep medicine community regarding the range of spindle frequency.

Recall that the proposed McSleep method detects spindles using a bandpass filter followed by the Teager operator. As a result, studies that may be interested in slow spindles (spindle frequency less than 13 Hz [67]) can set the passband as 10 Hz to 13 Hz or alternatively can set the passband as 13 Hz to 16 Hz for fast spindles. In this manner, the proposed method offers flexibility for the study of spindles, fast and slow alike. Moreover, since the computationally heavy transient separation algorithm needs to be run only once (for a fixed set of regularization parameters), the additional runtime in detecting slow and fast spindles separately is not significant.

Although the list of parameters required to be set by the user for the proposed method is not short, it is worth noting that only *λ*_2_ and *c* are the parameters that may need to be changed; all other parameters can be fixed for EEG signals that share the same sampling frequency. In Sec. 5.3 we explain how to tune parameters for large databases in a semi-supervised fashion.

### 4.2. Detection of global spindle activity

We illustrate the proposed McSleep method for detecting global spindle activity in a multichannel EEG. As described in Sec. 3.4, the Teager operator is applied to the channel mean of the bandpass filtered oscillatory component to detect spindles. We consider a sample segment of the multichannel EEG from the MASS database (Cohort 1, subset 2). For illustration purposes, we consider only three frontal, three central, three parietal and two occipital channels of the multichannel EEG. The EEG is shown in Fig. 8. Also shown in Fig. 8 is the channel mean of the bandpass-filtered oscillatory component, BPF(s), and its Teager envelope. Note that for visual clarity the channel mean of BPF(s) and its envelope are scaled. The detected spindles are shown using a binary vector at the bottom of Fig. 8. Figure 9 shows the spindles detected across the eleven channels from Fig. 8. Note that the detection is obtained by applying the Teager operator to each bandpass filtered oscillatory component and using the same threshold *c* for all the envelopes. Applying the Teager operator to the channel mean of BPF(s) has the effect of combining the detections from individual channels.

**Figure 8:**
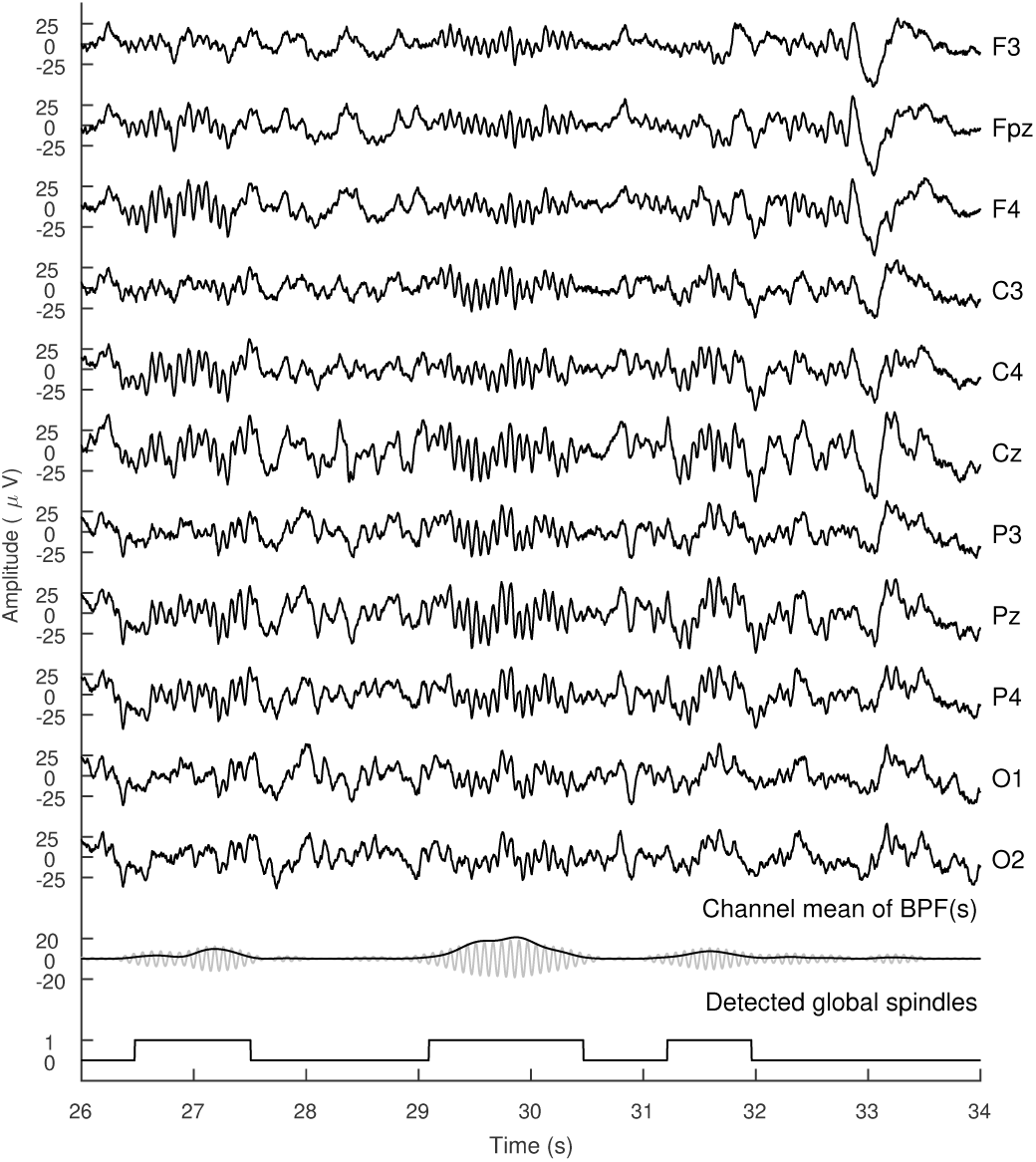
Global Sleep spindle detection on multichannel EEG using proposed McSleep method. Spindles that appear only on the left (alternatively only on the right) channels are also detected by the proposed method.

**Figure 9:**
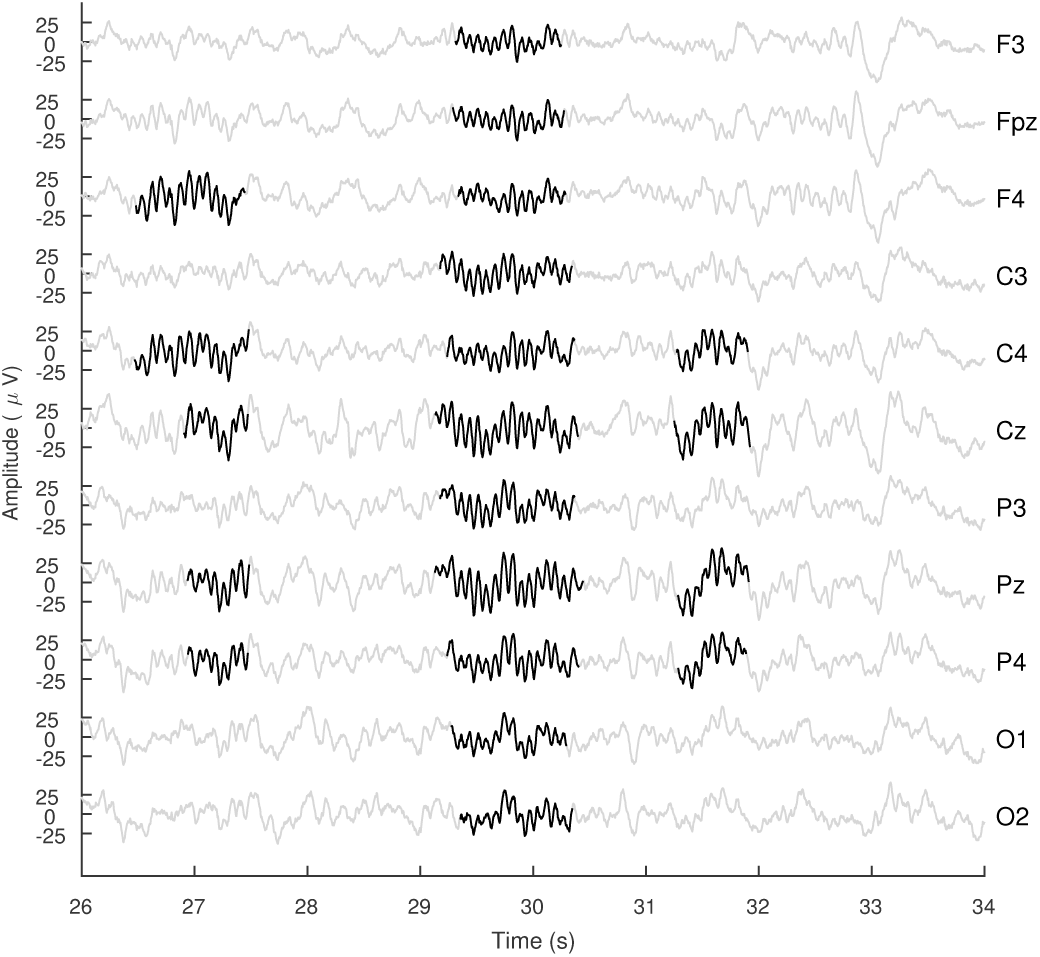
Sleep spindles detected (marked in bold) by applying the proposed McSleep detection method on multiple scalp channels of sleep EEG.

### 4.3. Comparison with existing single-channel detectors

We compare the proposed multichannel sleep (McSleep) spindle detection method with the following state-of-the-art automated detectors: Devuyst [22], Wendt [68], Martin [39], and DETOKS [50]. Note that for the proposed McSleep method, we apply the transient separation algorithm on three channels of the scalp EEG (FP1-A1, CZ-A1, O1-A1). As noted in the preceding subsection, post-estimation of the transient and oscillatory component, the Teager operator can be applied to each channel of the bandpass filtered oscillatory component to detect spindles in a single channel. The spindle detection carried out in this manner using the proposed method does not utilize fully the presence of multiple recorded EEG channels. Although, the proposed transient separation method certainly benefits using a multichannel EEG. In fact, the more channels present the better the separation between transients and oscillations for the input EEG.

Figure 10 shows the detection of sleep spindles for an example 3-channel EEG using the Devuyst, Wendt, Martin, DETOKS and the proposed McSleep methods. Also shown in Fig. 10 is the bandpass filter result for the different methods. Due to the absence of the implementation details for the bandpass filter used by Devuyst, we do not show the bandpass filter result in Fig. 10. Note that the experts have annotated three spindles at 357, 358 and 361 seconds visually for the central (CZ-A1) channel only.

**Figure 10:**
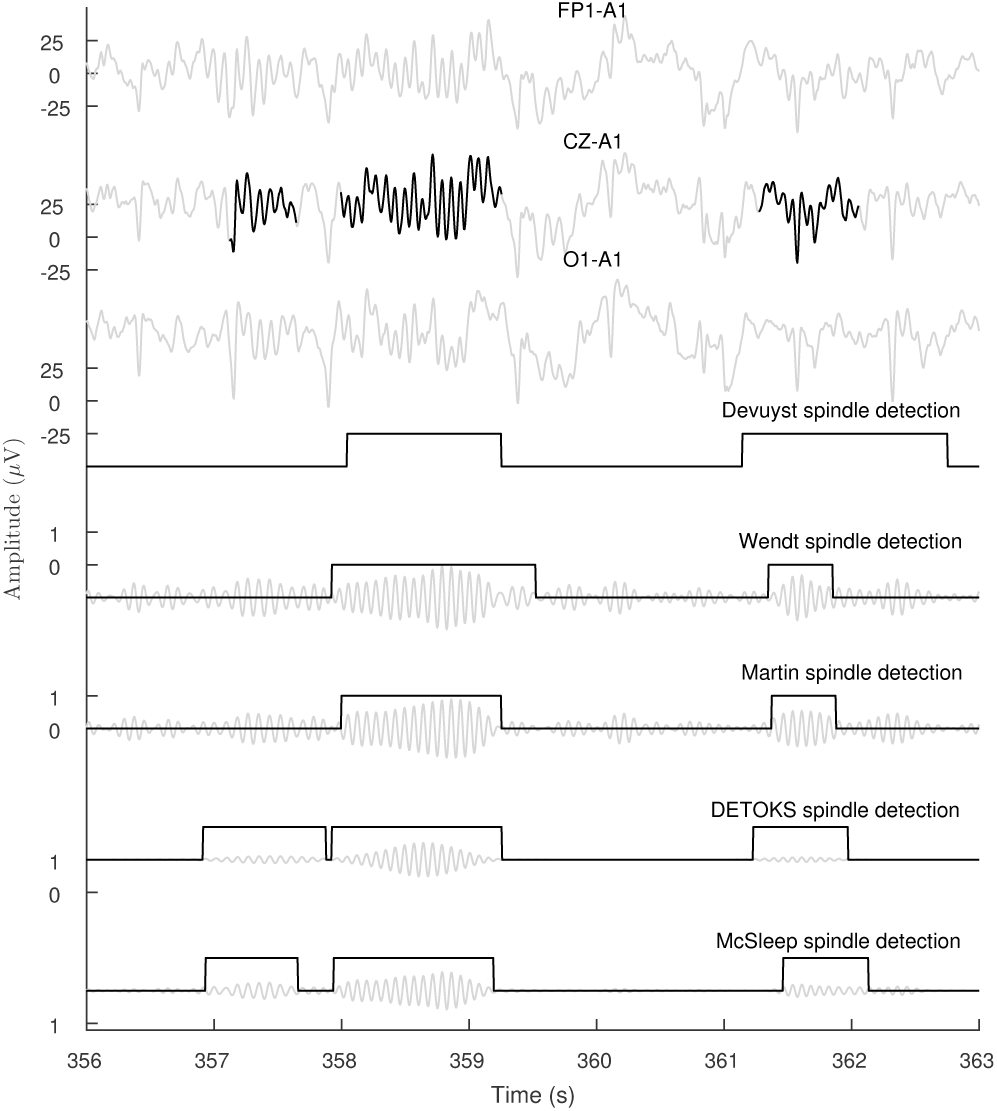
Comparison of proposed McSleep method for spindle detection with existing spindle detectors. Expert annotated sleep spindles are highlighted in black. Bandpass filter output is also shown in background for several methods.

The Devuyst, Wendt and Martin detection methods are not able to detect all the three spindles, with the Wendt method detecting a false positive spindle. The DETOKS method does detect all three spindles, but the estimated durations do not closely resemble the expert annotated spindle duration. Note that it is possible to increase the value of the Teager threshold for the DETOKS method to better match the duration of detected spindles. However, it is likely that this will discard previously detected spindles. On the other hand, the proposed McSleep method detects all three spindles and their estimated duration is similar to the expert detection. Moreover, the bandpass filter output for McSleep (only the central channel) shows the spindles much more prominently than the other methods.

Figure 11 shows the bandpass filtered EEG signal using the filter used by the Wendt algorithm [68]. As seen in Fig. 10, and reported in [50], the Wendt method detects false positive spindles due to the presence of transients. In particular, the transients in the sleep EEG excite the bandpass filter and as such the spindle activity does not appear prominent. This leads to the algorithm detecting false positive spindles in areas where non-oscillatory transients are present. It may also generally lead to a high number of false negatives. On the other hand, the proposed method seeks to first separate the transients and then use the estimated oscillatory component to detect spindles, thereby resulting in a much more prominent spindle activity in the bandpass filtered signal as seen in Fig. 12.

**Figure 11:**
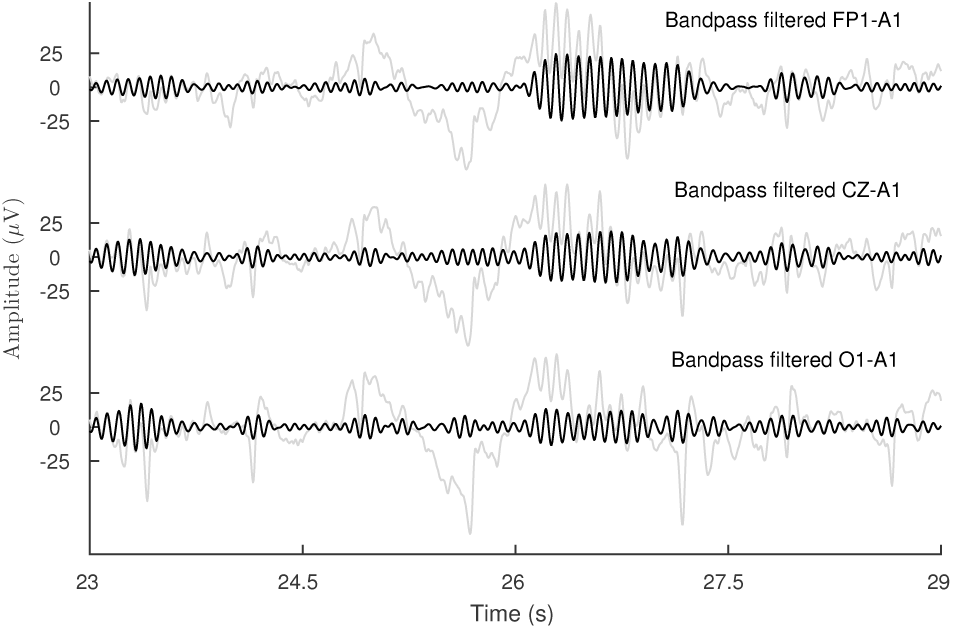
Bandpass filtered sleep EEG using Wendt algorithm [68]. The input EEG is shown in the background. As can be seen, the presence of transients excites the bandpass filter which may lead to false detections.

**Figure 12:**
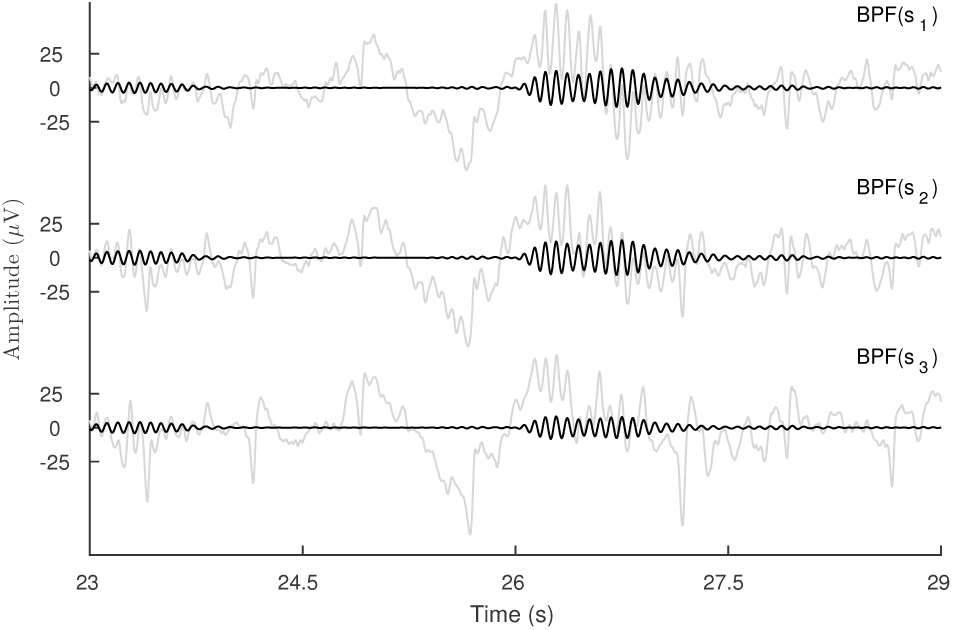
Bandpass filtered oscillatory component estimated using the proposed method. Due to separation of transients, the spindle activity is displayed more prominently.

## 5. Illustration of McSleep for single channel spindle detection

To further illustrate the proposed McSleep method we apply it to two online databases and compare the spindle detection results with several state-of-the-art automated single-channel detectors. Specifically we use the DREAMS [22] and the MASS [45] databases which have multichannel EEG recordings and spindles annotated only in the central channel.

### 5.1. Datasets and existing automated detectors

The DREAMS database^5^ was acquired using a 32-channel polygraph (BrainnetTM System of MEDATEC, Brussels, Belgium) [22]. The subjects possessed different pathologies (dysomnia, restless legs syndrome, insomnia and apnoea /hypopnoea syndrome) [23]. The online database provides 8 excerpts of 30 minutes from the whole-night recording. The excerpts contain three EEG channels (FP1-A1, C3-A1 or CZ-A1, and O1-A1), two EOG channels and one submental EMG channel. These excerpts were scored independently by two experts for sleep spindles. Out of the 8 excerpts, we use only 5 of the excerpts as these were scored by both the experts.

The MASS database [45] (Cohort 1, subset 2) contains full night multichannel EEG recordings from 19 healthy young subjects. The overnight recordings contain 19 scalp channels sampled at a frequency of 256 Hz. Of the 19 recorded channels, we use three frontal (F3, F4 and Fz), three central (C3, C4, and Cz) channels for the proposed McSleep method. All the recordings are annotated for spindles by two experts using the C3 channel and a linked-ear reference. The second expert annotated spindles using broad-band EEG signals (0.35 Hz - 35 Hz) and sigma filtered signals (11 Hz −17 Hz) similar to [53, 37]. Out of the 19 recordings only 15 were annotated by both the experts and as such we use those 15 recordings. We converted the visual annotations from the EDF+C format to a comma separated value (csv) file using the EDFBrowser software^6^

We use the following state-of-the-art automated detection algorithms: Wendt [68], Martin [39], Bodizs [8], Wamsley [66], Mölle [32], Devuyst [23], and DETOKS^7^ [50]. We refer the reader to [67, 49], and [50] for a review and the source code of the detection methods. We base the choice of detection algorithms for comparison on the availability of the source code for each of the algorithms. We apply the existing single-channel detectors to the central channel (C3 or Cz) of both the online databases. For the proposed McSleep method, we run the transient separation method on all the specified channels (3 for DREAMS and 6 for MASS) and apply the Teager operator to only the bandpass filtered central oscillatory channel for detecting spindles.

For both the databases the epochs containing electrode artifacts were identified visually, as in [24, 22], and discarded. The electrode artifacts considered were: lead movements and other body movement artifacts that result in abnormal jump in the amplitude of the EEG. Furthermore, we discard all detected spindles which are either less than 0.5 seconds or greater than 3 seconds [67].

### 5.2. Measure of performance

We use the expert detection (visually annotated spindles using the central channel) as a gold standard for evaluating the performance of the automated detection algorithms. We use the by-sample method of analysis, as described in [67] with the gold standard as the ‘union’ of the detections by both the experts. For the ‘by-sample’ rule, a time sample of the EEG is marked as a true positive (TP) if it was marked as a spindle by either of the experts and the automated detection algorithm. In this way, we calculate the true negative (TN), false positive (FP) and false negative (FN) values which lead to a 2 by 2 contingency table. These values are then used to evaluate the recall and precision scores of each of the detectors. Since spindle events are rare overnight, the specificity values will be abnormally high and may not lead to a proper analysis of the automated detector. The recall and precision scores are further used to calculate the F_1_ score, where the F_1_ score is defined as the harmonic mean of recall and precision Note that the F_1_ score ranges from 0 to 1, with 1 indicating a perfect detector. Similar to the F_1_ score, we also calculate Cohen’s κ [17] and Matthews Correlation Coefficient (MCC) [40].

### 5.3. Parameter Tuning

As described in Sec. 4.1 the illustration of the proposed McSleep spindle detection method on the online databases requires a set of optimal parameters to be chosen. Recall that the proposed McSleep method is a two-step detection process: first we estimate the transient and the oscillatory component and then use the oscillatory component to detect spindles by using a combination of bandpass filter and the Teager operator. The task-specific parameters such as the passband of the bandpass filter and block length are fixed to the same value as in Sec. 4.1.

Four key algorithmic-specific parameters are required to be set: *λ*_0_, *λ*_1_, *λ*_2_ and *c* (threshold for the Teager operator). The parameters *λ*_0_ and *λ*_1_ are set empirically to 0.3 and 6.5 for all the excerpts in the DREAMS database with a sampling of frequency of 200 Hz, whereas they are set as 0.6 and 7 for all the excerpts in the MASS database. For excerpts with a different sampling frequency in the DREAMS database, the values of *λ*_0_ and *λ*_1_ are scaled appropriately. The fixed value of *λ*_0_ and *λ*_1_ is chosen so as to ensure that the oscillatory component is relatively free of transient activity such as BCG or other cardiac artifacts. Note that we run only the transient separation algorithm, and not the entire proposed spindle detection method in order to fix *λ*_0_ and *λ*_1_.

Once the transient component is estimated the remaining parameters that need to be chosen are *λ*_2_ and *c*. For each subject, we choose a segment of the multichannel EEG (usually between 5-10 epochs) and run the proposed McSleep method (transient separation + spindle detection) for a grid of values of *λ*_2_ and *c*. We select the set of parameters that yield the highest F_1_ score for that particular segment. Figure 13 shows the F_1_ score as a function of *λ*_2_ and *c* for a 10 epoch segment for Excerpt 2 from the DREAMS database. It can be seen from Fig. 13 that the optimal set of parameters is *λ*_2_ *≈* 26.5 and *c* = 1.5. For the experiments that follow, the value of *λ*_2_ is varied in the range (25, 35) for the DREAMS database and in the range (45, 48) for the MASS database. The value of the threshold *c* is varied in the range (0.5, 3) for both the databases. Note that selecting the optimal parameters based on a small segment of the EEG may seem as over-fitting. As such, we suggest running the proposed method on the entire EEG a few times with parameters surrounding the optimal set obtained above.

**Figure 13:**
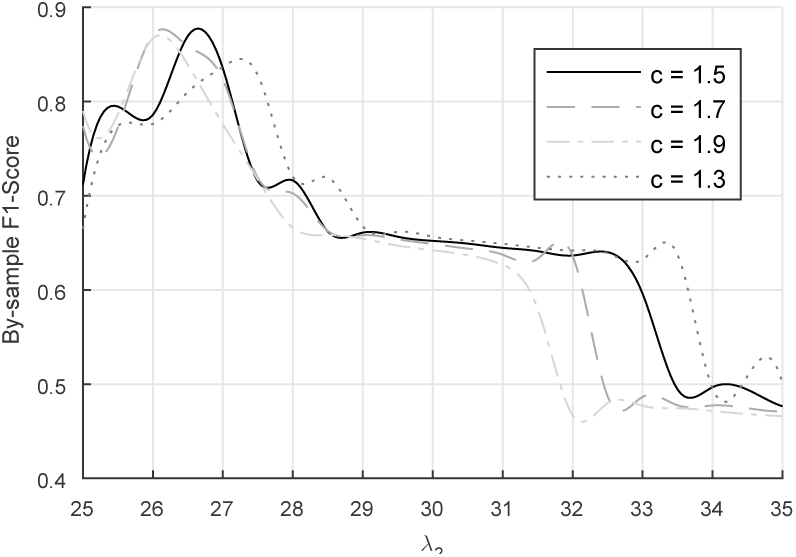
By-sample F_1_ score as a function of threshold *c* and *λ*_2_. For visibility only the *c* values that yield the highest F_1_ scores are shown. The test data is chosen from Excerpt 2 of the online database described in Sec. 5.

### 5.4. Results

The average F_1_, MCC, recall and precision values for the proposed McSleep method in comparison with the other existing detectors are listed in Table 1 for the DREAMS database and in Table 2 for the MASS database. Detailed statistical measures for the proposed method in comparison with the existing methods on both the databases are listed in Appendix 9.2.

**Table 1:**
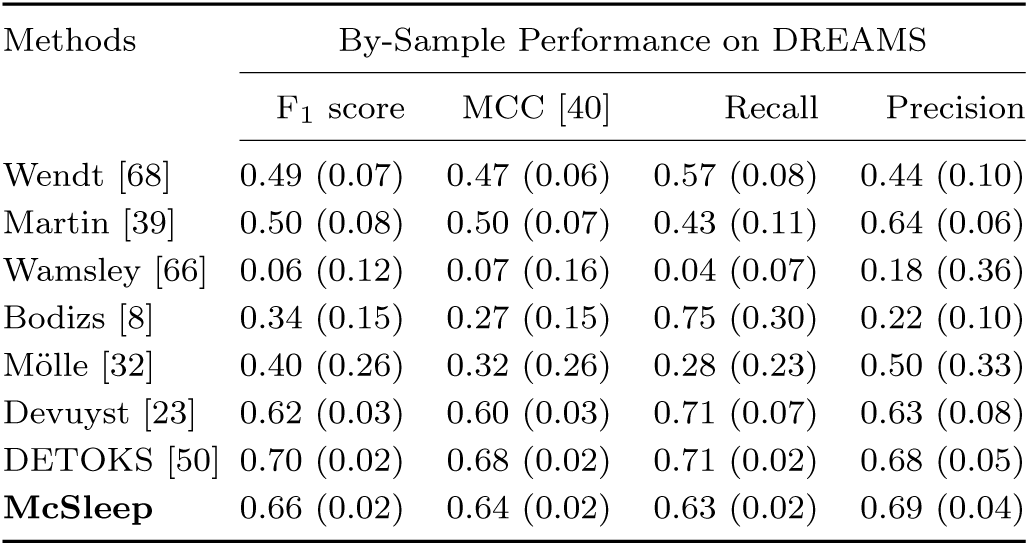
Evaluation of proposed method for sleep spindle detection as described in Sec. 5. Average values for the F_1_ score Matthews Correlation Coefficient (MCC), recall and precision over 5 excerpts are listed. Standard deviation values are shown in parenthesis.

**Table 2:** Average values of the F_1_ score, Matthews Correlation Coefficient (MCC), recall and precision over 15 EEG recordings are listed. The PSG recordings are from the MASS [45] database for which both the experts annotated spindles on the central channel (C3). (Standard deviation values are listed in parenthesis.)

| Methods | By-Sample Performance on MASS |  |  |  |
| --- | --- | --- | --- | --- |
| | $F_1$ score | MCC [40] | Recall | Precision |
| Wendt [68] | 0.54 (0.06) | 0.52 (0.05) | 0.55 (0.13) | 0.56 (0.12) |
| Martin [39] | 0.32 (0.20) | 0.32 (0.21) | 0.66 (0.33) | 0.29 (0.20) |
| DETOKS [50] | 0.60 (0.06) | 0.59 (0.05) | 0.55 (0.12) | 0.70 (0.11) |
| **McSleep** | 0.62 (0.06) | 0.60 (0.05) | 0.61 (0.09) | 0.64 (0.08) |

The proposed detection method on a 30 minute excerpt from the DREAMS database with three EEG channels (sampling frequency of 200 Hz) takes on an average 10 seconds on an Intel Core i7 cpu-based machine. This consists of 40 iterations of the transient separation algorithm, followed by bandpass filtering the oscillatory component and applying the Teager operator for spindle detection. The average runtime for the single channel detectors varied from 10 to 100 seconds with the most time taken by the Mölle detector [32]. For an overnight EEG recording (approx. 8 hours) with 6 channels the proposed method takes on an average 4 minutes to detect spindles in all the six channels. In comparison, a bandpass-filter-based single channel detection method takes on an average 1 to 5 minutes for detecting spindles in all 6 channels, while a transient separation based algorithm, such as DETOKS [50], takes on an average 8 minutes (run in parallel over all 30 second epochs of the six channels).

The proposed McSleep detection method can be run in two ways: either in parallel on 30 second epochs or on the entire overnight EEG (aprrox. 8 hours). We choose the former for the analysis presented in this paper. The epoch-by-epoch method of execution is done solely for faster runtimes. Moreover, it also enables the utility of the proposed method in an online mode^8^. The spindle detection is not affected whether the proposed method is run in parallel or on the entire overnight EEG. Furthermore, the proposed method can be run on user-chosen epochs as well.

### 5.5. Discussion

On the DREAMS database, the proposed McSleep detection method achieved better average F_1_ scores compared to other state-of-the-art detectors. The highest average F_1_ scores, however, were obtained by the DETOKS [50] method. The Martin [39] and Devuyst [22] methods performed relatively better than other detectors, in terms of the average F_1_ scores. On the MASS database, the proposed McSleep method achieved highest F_1_ scores. The DETOKS method [50] performed similarly but with slightly lower F_1_ scores (0.60±0.06 as compared to 0.62±0.06 for McSleep). However, for PSG 17, from the MASS database, the DETOKS method outperformed the proposed method. Note that the results for only the top four methods are shown in Table 2.

As shown in Sec. 4, the proposed McSleep method performs better when utilizing joint information from multiple EEG channels to separate the transients and oscillations. As a result, the performance of the proposed method is expected to, in terms of spindle detection against other existing detectors, increase proportionally with the number of EEG channels recorded (or provided online). For the DREAMS database, where only three EEG channels were provided, the proposed method didn’t outperform state-of-the-art single channel detectors (DETOKS in this case). On the other hand, for the MASS database, where six recorded EEG channels were used for McSleep, the performance of the proposed method is relatively better in comparison to other detectors. Furthermore, as noted in the preceding subsection, contrary to the belief that using multiple EEG channels might slow down an automated detector, the proposed McSleep method is faster than single channel methods run sequentially on multiple channels.

The transient separation algorithm for the proposed McSleep method and the one proposed in DETOKS [50] are quite similar. In particular, the regularization terms used for the transient component are same in both the methods, with the only difference being that the proposed method uses a multichannel input whereas DETOKS uses a single channel input. The notable difference between the McSleep and DETOKS method is in the use of the regularization term for the oscillatory component with the former using a low-rank regularization and the latter using a sparse STFT regularization. For the case of a single channel input, the proposed objective function in (18) penalizes sum of absolute values of overlapping blocks. In comparison, DETOKS penalizes sum of absolute values of overlapping STFT blocks. As such, the proposed McSleep method and DETOKS are expected to perform similarly, especially for a dataset that either has EEG channels only from one hemisphere of the brain or where the experts viewed only the central channel on the screen while scoring spindles.

As noted in the Sec. 1.1, a simple method for detecting sleep spindles across all the channels of scalp EEG is to run the existing single channel detectors sequentially channel-by-channel and if required, combine the resulting detections. In order to combine the single channel detections a majority-vote type method may be used. However, ignoring detected spindles on the basis that they do not pass a majority vote can possibly lead to a high type II error, especially for studies that are investigating the spatial distribution of sleep spindles overnight (for e.g., see [9], [51]and the references therein). Another method, perhaps simpler than majority voting, for combining the detection is to use a ‘union’ rule: a sleep spindle detected in any one channel is a valid detected spindle. However, such a union rule generally leads to high type I error (high false positives are reported).

Consider the EEG segment shown in Fig. 14, where the expert annotated spindle is at 109.3 seconds. Figure 14 shows the spindle detection obtained using three methods: DETOKS run on only the central EEG channel, DETOKS run on all three channels sequentially with separate parameter tuning, and the proposed McSleep method. The single channel DETOKS method run on the central channel alone detects a false positive. Whereas, using the union rule, the all-channel DETOKS detects the true positive spindle but also retains the false positive spindle. In contrast, the proposed method, due to a better separation of transients and oscillations, does not detect the false positive spindle while correctly detecting the expert annotated spindle. Similar behavior is observed in another EEG segment shown in Fig. 15.

**Figure 14:**
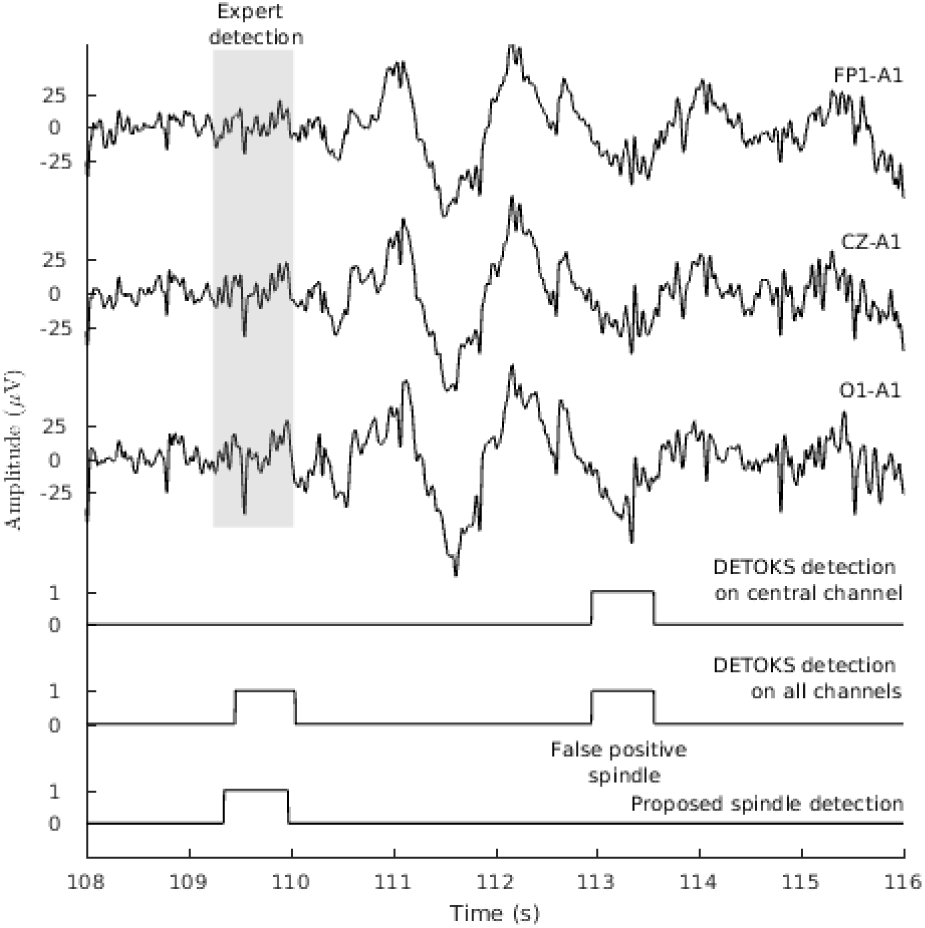
Comparison of the proposed McSleep detection method and the DETOKS detection method. The DETOKS method detects a false positive when run on either a single central channel or on all the channels. The EEG is obtained from excerpt 2 of the online EEG database [23].

**Figure 15:**
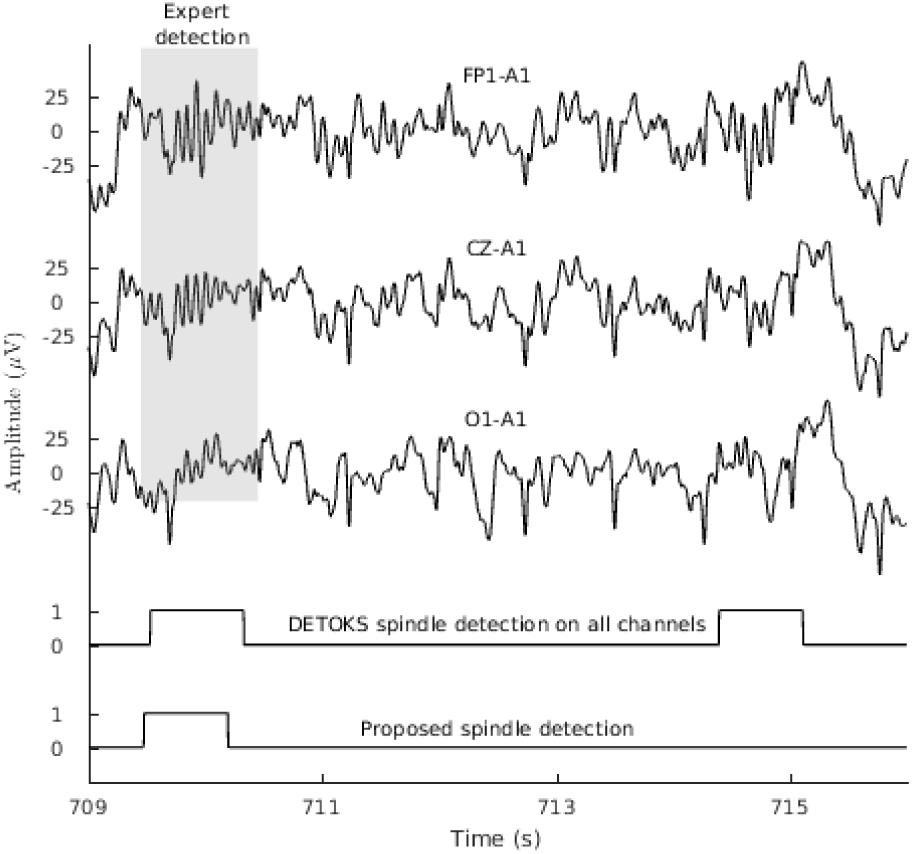
Proposed McSleep detection in comparison with DETOKS method run sequentially on all channels. The proposed detection contains fewer false positive spindles due to better separation of transients.

## 6. Conclusion

Automated spindle detectors that consider only a single channel of the recorded EEG are blind to the presence of spindles in other channels. On the other hand, a human expert annotating spindles visually based on the AASM manual may be subliminally influenced by the presence of spindles in other EEG channels visible on the screen. In order to mimic such a human behavior and simultaneously utilize joint information from multiple EEG channels, we propose a multichannel sleep spindle detector using a non-linear signal model for the multichannel EEG. The proposed spindle method uses a multichannel transient separation algorithm that separates the non-rhythmic transients from spindle-like oscillations, thereby estimating the components of the proposed non-linear signal model. The transient component is modeled as a piece-wise constant signal with a baseline of zero whereas fixed-length blocks of the multichannel oscillatory component are assumed to be low-rank. The oscillatory component is then used to detect spindles. A fourth order Butterworth bandpass filter and the Teager operator are used to detect spindles following the transient separation process.

Several examples are shown to illustrate the utility of the proposed multichannel spindle detector and a comparison with state-of-the-art single channel spindle detectors is performed using two publicly available online databases. A fast run-time and better average F_1_ scores enable the proposed multichannel sleep spindle detector to be a valuable tool for studying the architecture of sleep spindles and tracking their behavior in sleep EEG. While the degree to which a human expert is influenced by the presence of spindles in channels visible to him on the screen is an open question, the proposed method shows that using multichannel EEG certainly yields better estimation of transients and spindles alike. Thereby, increasing the agreement between human expert and an automated algorithm for spindle detection.

## 7. Acknowledgments

The authors thank the anonymous reviewers for their detailed suggestions which have improved the manuscript considerably.

## 9. Appendix

### 9.1. Solution to the Least-Squares step of the proposed transient separation algorithm

We derive the solution to the least-squares sub-problem in (21a), which is written below for clarity to the reader.

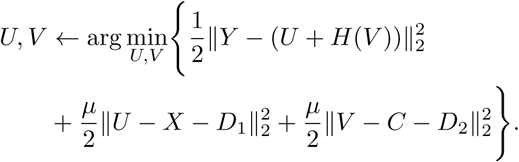

We make the following substitutions

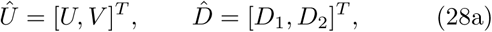

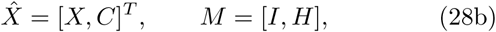

and re-write the objective function as

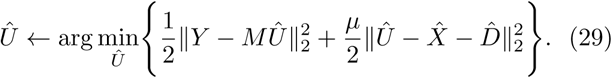

The solution to (29) can be written explicitly as

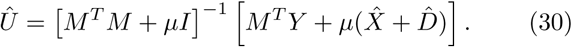

The inverse in (30) results in inverting a dense matrix consisting of *M* ^*T*^ *M*. In order to efficiently compute the explicit solution, we use the Matrix Inverse lemma [70, 64]. As such, the solution in (30) can be written as

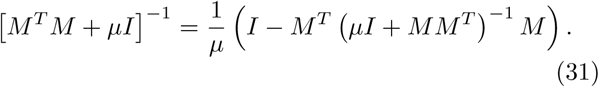

Note that the operator *H*, where *H* = Φ^*T*^, is implemented in this paper for perfect reconstruction. Hence, we have

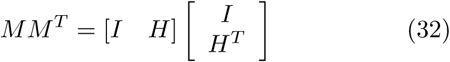

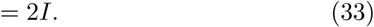

As a result, the inverse in (30) can be written as

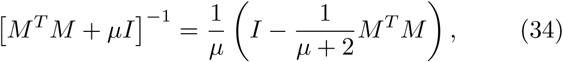

which leads to the following explicit solution for 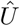,

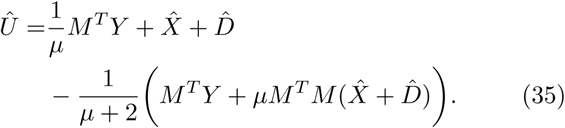

Combining (35) and (28), we get the following steps for obtaining the solution to the objective function in (21a).

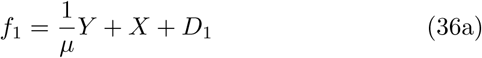

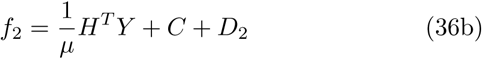

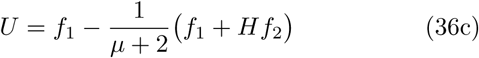

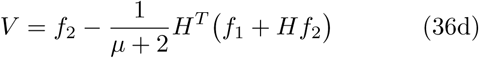

Note that *H*^*T*^ *Y* can be pre-computed outside the iterative loop for speed.

### 9.2. Illustration of McSleep for sleep spindle detection

We illustrate the utility of the proposed multichannel spindle detection (McSleep) method by applying it to detect spindles on two online databases: the DREAMS database [22] and the MASS dataset [45]. We also run several state-of-the-art spindle detectors and compare their performance with McSleep. For a summary of each of the measures of performance, we refer the reader to [67].

**Figure 16:**
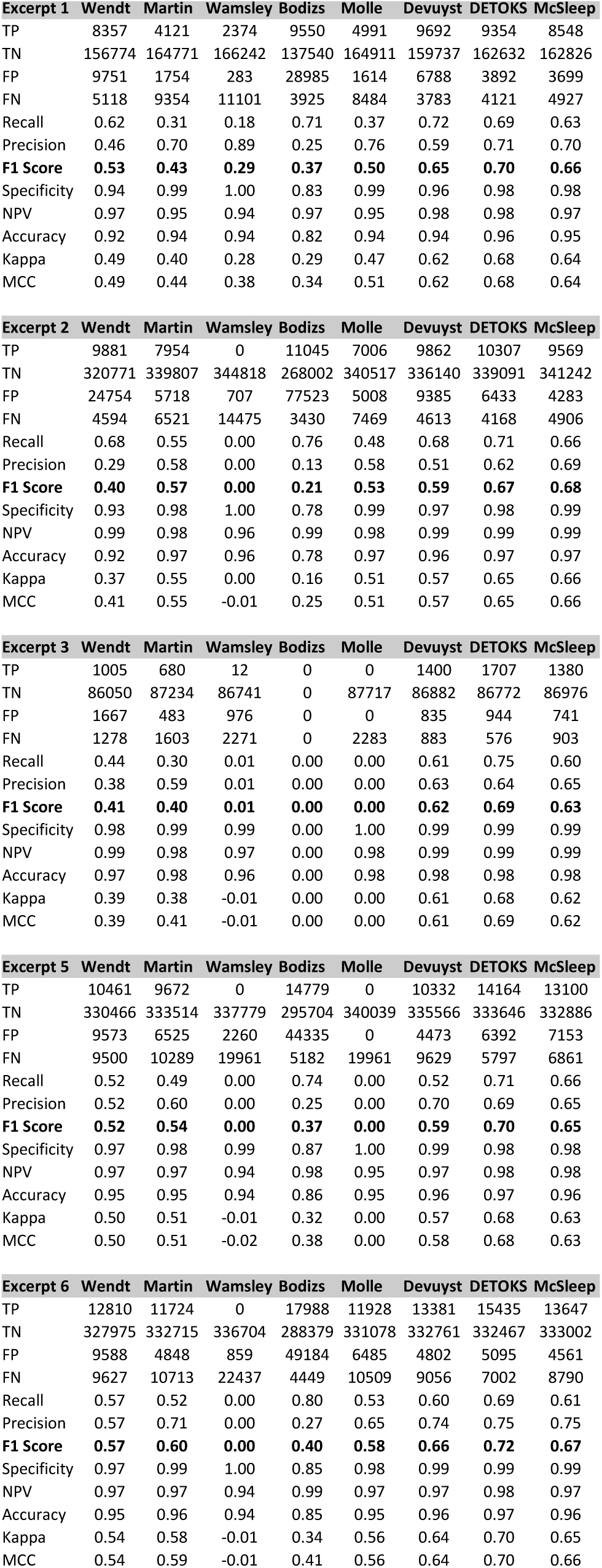
Performance of proposed McSleep and existing sleep spindle detectors on the DREAMS database.

**Figure 17:**
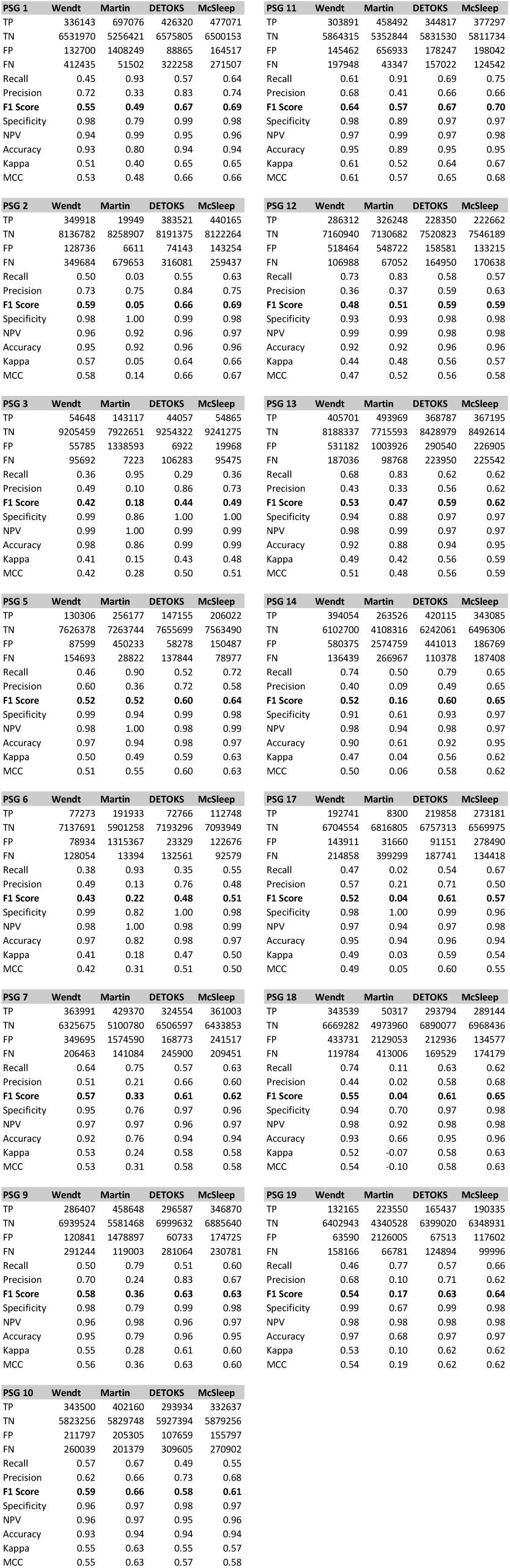
Performance of proposed McSleep method and existing sleep spindle detectors on the MASS database. (The numbers are best viewed on the pdf file by zooming in.)

## Footnotes

1 Given an input matrix, a low-rank approximation aims to find a matrix similar to the input matrix but with a reduced rank.

2 Note that using a generic amount of overlap does not guarantee perfect reconstruction i.e., Φ^*T*^ Φ ≠ *I*.

3 https://github.com/aparek/mcsleep.git

4 University of MONS - TCTS Laboratory (S. Devuyst, T. Dutoit) and Universite Libre de Bruxelles - CHU de Charleroi Sleep Laboratory (M. Kerkhofs)

5 http://www.tcts.fpms.ac.be/∼devuyst/#Databases

6 http://www.teuniz.net/edfbrowser/

7 https://github.com/aparek/detoks

8 An online algorithm is able to process the input data in a piece-by-piece fashion without requiring the presence of an entire signal.

